# Modelling the decline in *Sporobolus anglicus* detections toward functional eradication: a case study in the Marlborough Sounds, New Zealand

**DOI:** 10.64898/2025.11.30.691427

**Authors:** Rae Lerew, Christoph D. Matthaei, Stephanie S. Godfrey

## Abstract

**Summary:** *Sporobolus anglicus* (C.E. Hubb) P.M. Peterson & Saarela (synonym *Spartina anglica*) is a highly invasive coastal weed that forms dense monocultures in intertidal mudflats and estuaries, displacing native reedbeds and associated fauna. Introduced to Aotearoa New Zealand from Britain in the early 1900s to aid coastal land reclamation, it became a conservation concern by the 1960s, prompting control efforts in the South Island from the 1970s.

This study presents a detailed *S. anglicus* eradication case study from the Marlborough Sounds, where detections are nearing zero. We model changes in detection probability over time in Te Hoiere / Pelorus Sound, a complex estuarine catchment. We aim to inform control efforts and assess the effectiveness of ongoing management, by evaluating the probability of non-detection as a proxy for functional eradication.

Using data from the Department of Conservation (DOC), we analysed detections across 79 search blocks between 2013 and 2024. A generalised linear mixed model was used to generate predicted detections through to 2040, using a modified dataset with pseudo-zero values for probable absences. The data were modelled as presence/absence with a binomial distribution, to identify the first year with a <0.01 probability of a positive detection (upper 99% confidence limit). Results suggest that by 2032, the likelihood of further detections under current management practices is remote, and that functional eradication may have occurred. We interpret this decline in detection probability as indicative of management success.

Model outputs can support decision-making as to when active surveillance might reasonably be ceased. To accelerate the tail-end of eradication efforts, we recommend intensifying search effort and widening delimitations within the catchment over the next five years, to ensure removal of any remaining individuals. We also propose the use of environmental DNA as a cost-effective backstop for after operational wind-down.

**Implications for Managers:**

- Our modelling predicts that *S. anglicus* detections in Te Hoiere / Pelorus Sound should decline to levels consistent with functional eradication by 2032, and may render continued manual surveillance uneconomical after this date.
- Full, regular and repeated surveillance of suitable habitats is needed within a five-year intensive monitoring period at the tail-end phase of eradication, as detections approach zero and managers consider withdrawal.
- While statistical analyses support eradication management decisions, absolute certainty of absence is unattainable. Decisions must balance technical feasibility with practical risk tolerance.
- Environmental DNA could provide an effective post-withdrawal monitoring tool, to allay the risk of re-invasion.

## Introduction

Cordgrasses of the genus *Sporobolus* (C.E.Hubb) P.M.Peterson & Saarela (synonym *Spartina*) are highly invasive in coastal areas globally, including in Aotearoa New Zealand, where they are classified as environmental weeds posing significant threat to native ecosystems (McAlpine & Howell, 2024; Wang et al., 2023). *Sporobolus* species rapidly occupy intertidal mudflats, often completely displacing native reedbeds, causing sediment accretion and the reclamation of estuarine environments (Lee & Partridge, 1983; Strong & Ayres, 2013; Sheehan & Ellison, 2014). Their global impacts on saltmarshes include lowering fish species’ abundance and diversity (Harrison-Day et al., 2023), reducing benthic invertebrate communities (Lu et al., 2022; Xu et al., 2024) and impacting the foraging success and habitats of waterfowl (Lyu et al., 2023; Xu et al., 2024). These cordgrasses were purposefully introduced to Aotearoa in the early 1900s, to increase flat land for industrial development and agricultural use (Lee & Partridge, 1983). Due to their ecological impacts, they are now the focus of eradication efforts, and the genus is one of twelve weeds included in Aotearoa’s “war on weeds” (Department of Conservation, 2016).

While much of what is known about *Sporobolus anglicus* (C.E. Hubb) P.M. Peterson & Saarela (synonym *Spartina anglica*) stems from Northern Hemisphere contexts, few studies examine its control and eradication in the Southern Hemisphere (Ainouche & Gray, 2016; Harrison-Day et al., 2023). In Australia, research has focussed on ecological impacts (Hedge & Kriwoken, 2001; Shimeta et al., 2016) and post-removal erosion (Sheehan & Ellison, 2015; Sheehan, 2008), but offers limited insight into long-term management outcomes (Kriwoken & Hedge, 2000). Internationally, work from America and Europe highlights persistent challenges to eradication, including re-invasion risk from missed control years (Reeder & Hacker, 2004), imperfect detectability (Patten et al., 2017), and the importance of sustained control efforts backed by political will (Hammond & Cooper, 2002; Milne, 2008). China’s experience with *Sporobolus alterniflora,* which invaded 70% of the coastline in just fifty years, triggered a national eradication campaign in 2022, underscoring the weed’s invasiveness (Xu et al., 2024). However, biological variation between regions, such as reproductive traits, dispersal, and seedbank persistence, limits the applicability of these insights to Aotearoa (Xiao et al., 2010). While these global studies affirm the value of local eradication efforts, they also signal persistent obstacles including detectability, reinvasion, and the need for consistent monitoring.

Given these challenges, projecting a possible timeline toward a functional eradication threshold is essential for effective management planning (Kerr et al., 2016). Because absolute absence of a weed is rarely verifiable in ecological systems, ‘functional eradication’ (henceforth ‘eradication’) serves as a more pragmatic goal, defined here as the point where the likelihood of future detections is low enough that active surveillance can be justifiably halted (Rout, 2017; Green & Grosholz, 2020). As detection rates decline, managers must therefore be equipped to adapt thresholds of eradication – and their expectations (Rejmánek & Pitcairn, 2002). While a plant’s biological traits are undoubtedly pivotal in deciding whether complete extirpation is possible, eradication can only be achieved where sustained funding and institutional will can outlast biological thresholds for re-invasion (Simberloff, 2003; Regan et al., 2006; Panetta 2015).

Biologically, *S. anglicus* eradication appears feasible. Its seeds remain viable for a maximum of four years and have a germination-to-seeding period of two to three years (Gray & Benham, 1990; Partridge, 1987; Hubbard 1970). Nonetheless, prolific seed and dynamic tidal spread increase re-invasion risk and complicate timelines (Mullins & Marks, 1987). Economically, project managers must decide how best to direct surveillance efforts as detections approach zero – and ultimately, decide when to stop looking (Ramsey et al., 2023). In Aotearoa, ad-hoc assessments have commonly been used to determine exit strategies rather than quantitative tools, resulting in protracted projects and few confirmed successes (Howell, 2012; Regan et al., 2006).

To improve outcomes, eradication programs benefit from predictive models that estimate future detection probabilities using surveillance data. These tools help managers balance the cost of continued surveillance against the environmental cost of premature project withdrawal and the chance of reinvasion (Regan et al., 2006). However, most existing models rely on plant detectability estimates to account for false negatives (the probability that a plant goes undetected) due to observer bias in the field (Ramsey et al., 2023; Moore et al., 2011). While detectability is commonly estimated through occupancy modelling using repeat-survey data (MacKenzie et al., 2002) or controlled re-survey trials (Ramsey et al., 2023), both are data and resource intensive. Occupancy modelling may not be suitable for near-eradication datasets, where numerous records of absence provide little contrast in the data with which to estimate detectability (Sileshi et al., 2009). The gold-standard of detectability assessment, controlled re-surveys, may be impractical and economically out of reach for many management programs (Moore et al., 2011; Ramsey et al., 2023; West & Havell, 2019). Furthermore, detectability varies by species, site, method and season, requiring the re-estimation of parameters for each situation (Chen et al., 2009; Wintle et al., 2009; Ramsey et al., 2023).

These factors underscore the need for modelling approaches that work with available ‘real-world’ detection data. In Aotearoa, weed eradication attempts often extend for decades without resolution, and many rely on qualitative scenario planning (Howell, 2012; Hulme, 2012). If limitations are clearly acknowledged, detection-data models could help managers plan exit strategies from weed control projects. In this study, we use a *S. anglicus* eradication project in Aotearoa to model detection trends and estimate a probable timeline to functional eradication.

### Sporobolus anglicus: a case study in the Marlborough Sounds

The common cordgrass *S. anglicus* is a vigorous hybrid between *S. maritima* and *S. x townsendii* (Muhammad & Ki, 2024), first introduced near Havelock Estuary in the Marlborough Sounds in 1952 to reclaim land for a new shipping port (Millen & Waddel, 2003). It was part of a series of deliberate *Sporobolus* introductions around Aotearoa, and continued to spread via natural dispersal (Partridge, 1987).

By the 1960s, awareness of its ecological impact had increased, as estuarine values were threatened against a backdrop of rapidly expanding infestations (Partridge, 1987; Millen & Waddel, 2003). This came to a head in 2013, when the NZ Department of Conservation (DOC) assessed the feasibility of *Sporobolus* eradication in the South Island (Brown & Raal, 2013). The report concluded that eradication was achievable under certain conditions, including centrally coordinated management, sufficient resourcing, consistent methods of control and survey, and “intensive and extensive surveillance” to confirm eradication success (Brown & Raal, 2013).

The Marlborough Sounds management program now exemplifies many of these conditions, with Pelorus Sound / Te Hoiere the focus of initial *S. anglicus* infestations, and subsequent control efforts. Te Hoiere is a single estuarine system with extensive intertidal reedbeds at the heads of bays and inlets (Urlich & Handley, 2020). Its three largest estuaries are Kaituna, Mahakipawa and Kaiuma, summing approximately 1025 ha and bordered by 560 km of coastline (Bowie, 1963). By 2004, *S. anglicus* had formed dense monocultures in these landward estuaries, displacing native reeds and rushes such as *Apodasmia similis* and *Juncus krausii* (Figure 1). The system’s complexity and large size make Te Hoiere a fascinating case study for understanding *Sporobolus* eradication and spread (Figure 2).

**Figure 1:**
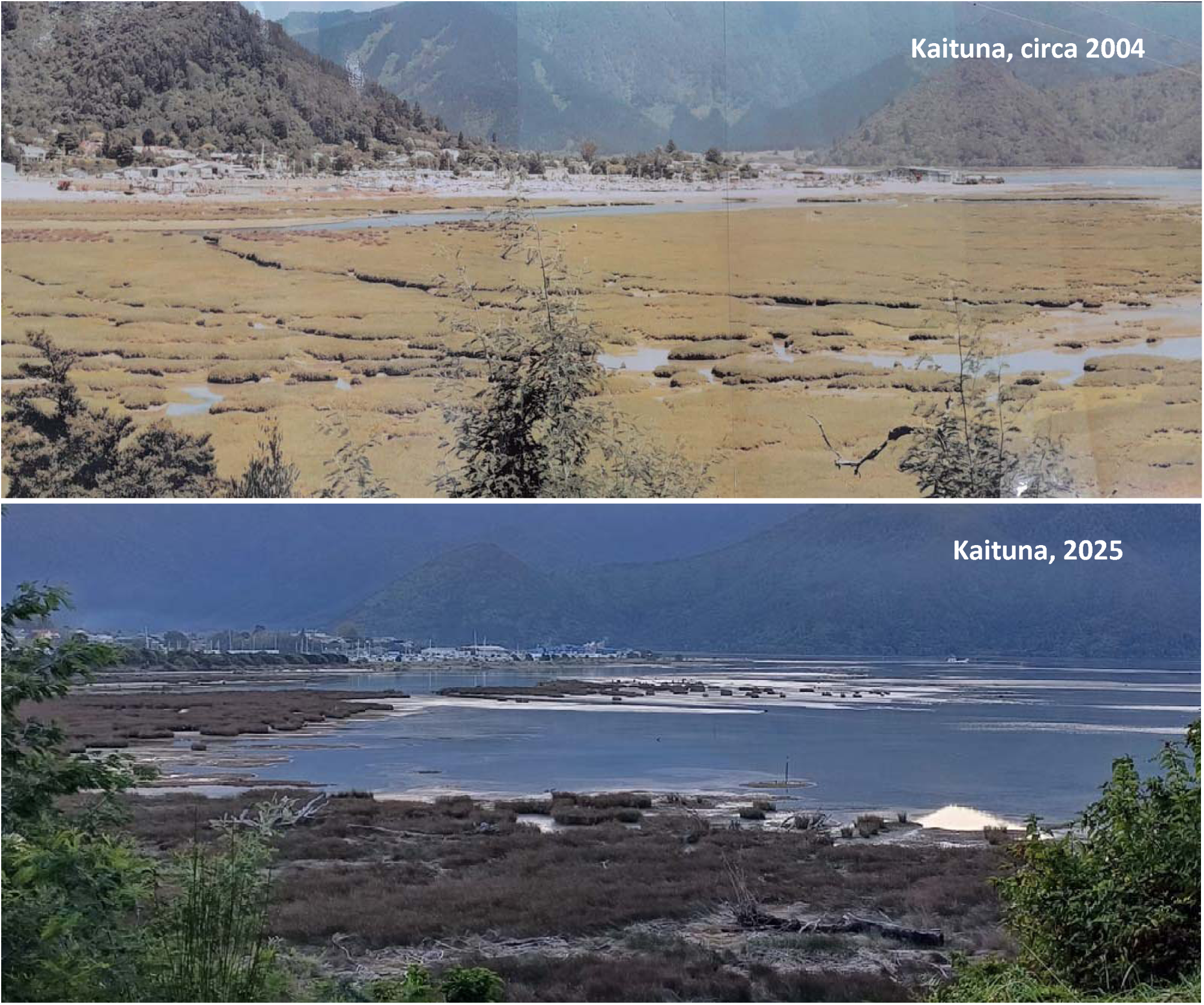
The top photograph was taken prior to the initial herbicide knockdown in 2004/2005 (exact date and photographer unknown). It shows dense swards of *S. anglicus* in Kaituna Estuary, looking across to the Havelock township from Queen Charlotte Drive. Almost all indigenous reedbeds have been displaced, barring isolated patches of reeds (seen in brown, left-hand). The bottom photograph was taken from a comparable location in early 2025, showing an absence of *S. anglicus*, and the re-growth of native reedbeds (image courtesy of Clinton Wadsworth).

**Figure 2:**
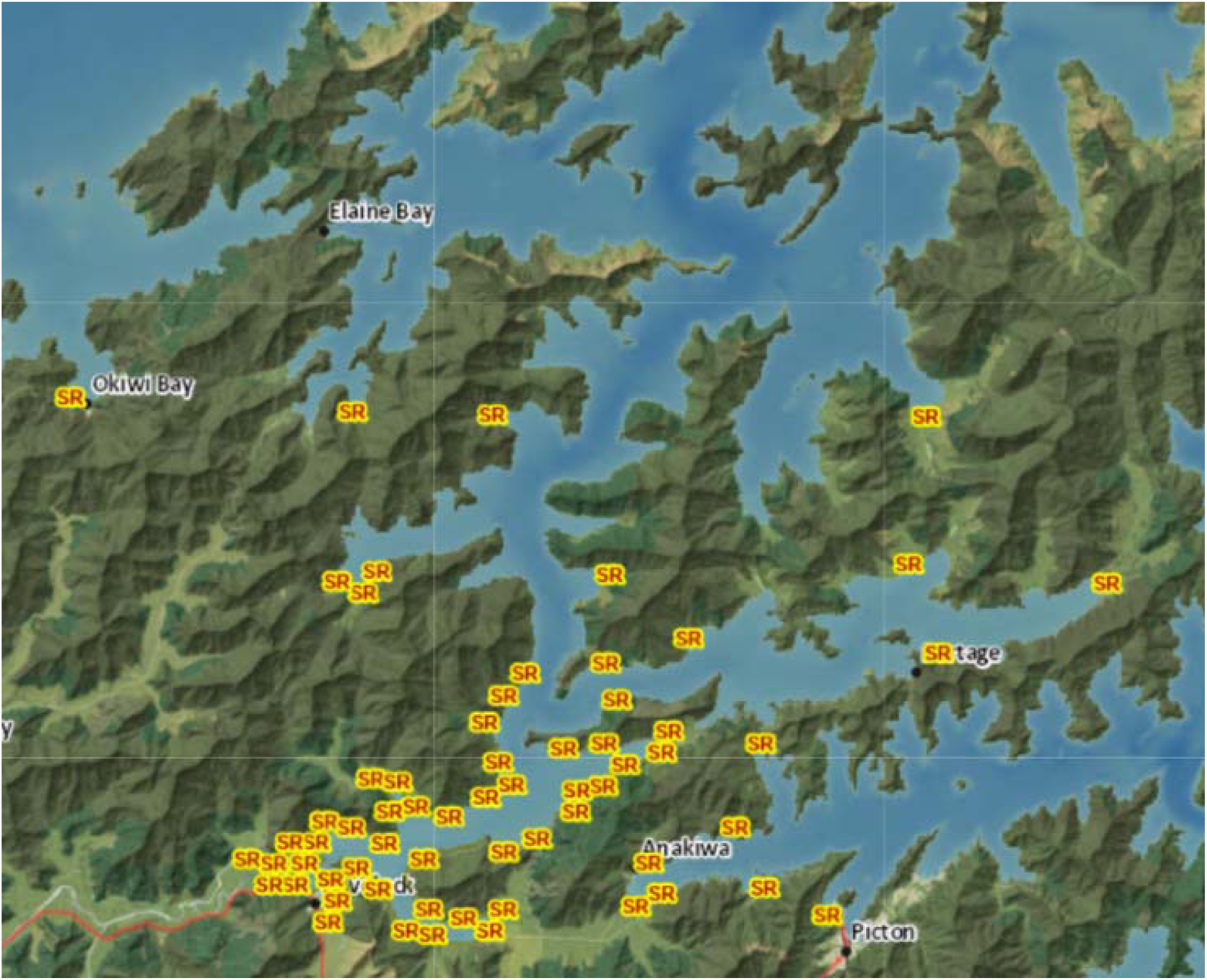
Map of cumulative *S. anglicus* detection hotspots in the Marlborough Sounds, with innermost Te Hoiere showing the densest detections. These are centred on the ‘Kaituna’, ‘Mahakipawa’ and ‘Kaiuma’ estuaries near Havelock township, seen bottom left. Source: Smartmaps.marlborough.govt.nz, ‘*Biosecurity Pest Plants’*, retrieved 28 May 2024.

Initial knockdown control in Te Hoiere began in 2004, with helicopters and tankers deploying thousands of litres of herbicide (P. Clerke, pers. comm., May 17, 2024). In subsequent years, teams have returned annually from January to March, to detect and remove any remaining plants before seed has set (Brown & Raal, 2013; D. Palmer, pers. comm., May 2024). Detected plants are now controlled using Haloxyfop-P-methyl ester as a spot spray at 1.5 g L ¹ (active ingredient) (C. Wadsworth, pers. comm., May 2024). Herbicide use for *Sporobolus* control has little negative impact on saltmarsh ecosystems (Wang et al., 2023).

Following early knockdown and a sharp decline in detections (Table S1), the need for a detailed surveillance framework was identified to track progressively scant detections. In 2013, Te Hoiere was divided into search blocks - defined survey polygons of varying sizes, using natural boundaries such as stream banks and roadsides (C. Wadsworth, pers. comm., May 2024). These search blocks form the basis of our study dataset, with *S. anglicus* detections within each block recorded for every year of treatment. A detection is defined as “an individual tiller or patch at least 10 m from another individual tiller or patch” (Brown & Raal, 2013). A team of four people grid-search each block shoulder-to-shoulder, ensuring complete coverage (Figure 3).

**Figure 3:**
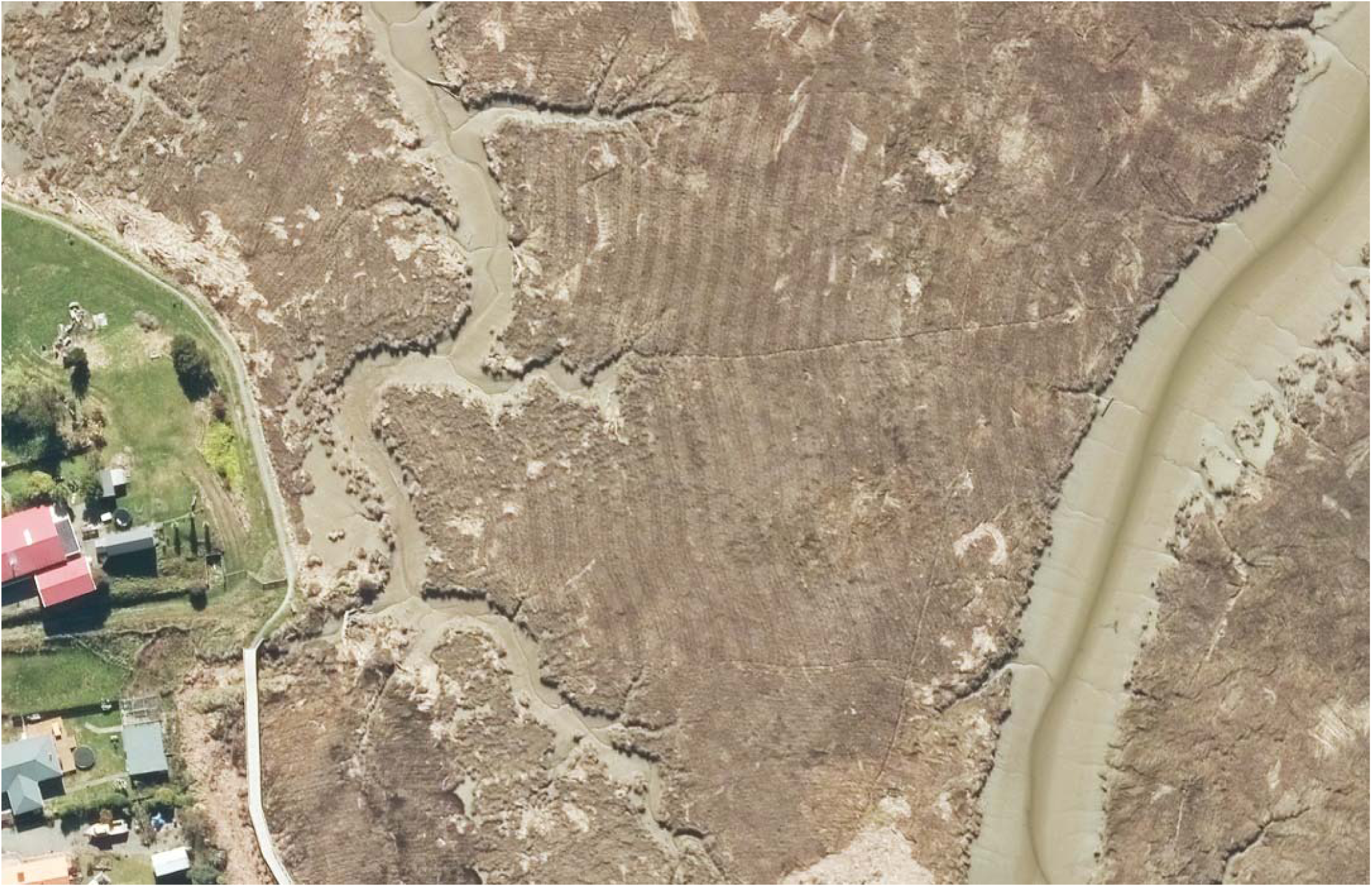
Aerial image of native reedbeds within search area ‘Kaituna’, adjacent to Havelock township, showing the consistency of the grid-search survey method. Pale and darker stripes represent different directions of travel as workers comb back and forward through the reeds. Within each directional stripe, the individual walking paths of four workers can be seen. Source: *Marlborough 0.25m Rural Aerial Photos (2023–2024)*, licensed under CC BY 4.0. Retrieved from LINZ Data Service, May 2025: https://data.linz.govt.nz/layer/114135

Despite the thoroughness of this search method, observer bias and inattention can still result in plants being overlooked (Chen et al. 2009; Moore et al. 2011), and management must balance funding constraints and seasonal accessibility with the need for sustained detection pressure. Annual cordgrass detections have now plateaued at numbers near-to-zero across search blocks, making surveillance efforts increasingly difficult to direct (D. Palmer, pers. comm., May 2024). Additionally, plants are occasionally detected in unexpected areas, apparently unrelated to historic detections, possibly due to tidal movement or buried plant matter (Strong & Ayres, 2013). These factors are complicated by the convoluted waterways, where potential habitat is too large to search entirely year-on-year.

Effective weed management requires a clear understanding of a species’ spread and the effectiveness of ongoing search and removal efforts (Simberloff, 2003; Hulme, 2020). At this stage of the program, detection modelling may offer a way forward. By estimating future *S. anglicus* detection probabilities in Te Hoiere, we can assess the usefulness of continued surveillance, and inform when functional eradication might reasonably be declared. We seek to address the following aims, based on model results:

1. Model *S. anglicus* detections in the Marlborough Sounds to estimate the probability of detecting plants in future survey years.
2. Use changes in detection probability over time to infer the likely trajectory toward functional *S. anglicus* eradication.
3. Use model outputs to evaluate the success of the management program, in the context of practical eradication thresholds.

## Methods

We obtained data on *S. anglicus* detections in Te Hoiere from DOC, Marlborough Sounds District Area Office. The dataset includes the number of *S. anglicus* detections and search effort over 12 seasons, from 2013 to 2024, for each of 79 separate search blocks. Because search effort was proportional to block size due to consistent search methods, detections were expected to scale with block size. We therefore did not standardize block size in our analysis. Search effort for blocks ranges from 0.25 to 4 days, corresponding approximately to the area of each block.

Given that every search block is not visited every control season, there are many missing values that may reduce model precision. The decision to not visit a block in a given year is driven by logistical considerations and prior detections. Blocks with repeated absences in prior years are considered less likely to record detections, and those blocks may be temporarily omitted from annual surveillance. Regardless, all blocks remain on an intermittent rotation to ensure that even those with repeated past absences are regularly visited.

To improve the accuracy of our models given these factors, we created a dataset with pseudo-zero values in place of selected missing values, in cases where near certainty of *S. anglicus* absence could be reasonably determined. We used the presence/absence pseudo-zero dataset in our model to generate more realistic predictions of eradication years, with high confidence.

The pseudo-zero dataset was created by inserting zeros where multiple searches in previous or subsequent years clearly indicated that a record of absence was highly likely for that year. For example, in 2015, 2018 and 2020, the ‘Matai-Tawa Shores’ block was surveyed with no detections recorded. Although it was not visited between those years, it is highly likely that in 2016, 2017 and 2019 cordgrass would not have been detected had the block been searched, even accounting for human error, given the three records of absence. We assessed the entire dataset in this way and replaced missing values with zero-values where appropriate. In some cases, missing values were retained. For example, ‘Cemetery’ block recorded records of absence for 2018 to 2020. It was not searched in 2021, but a plant was found in 2022. We therefore retained the missing value for 2021, as the 2022 find may well have been present in 2021, despite many records of absence prior to that year.

### Study Design

We divided the 79 search blocks into 13 search areas for analysis, grouped by geographic proximity (Figure S1). Most search blocks fall into distinct areas such as an inlet or estuary, or on either side of a main river channel. The 13 search areas allowed us to model *S. anglicus* detections in geographically distinct areas of Te Hoiere, whilst improving model convergence. The true data with missing values omitted came to a total of 500 observations, whereas the equivalent pseudo-zero data had a total of 907 observations. GPS points were taken for each detection, and detections were mapped by both DOC and MDC on their respective GIS systems.

### Statistical Analysis

We used the statistical software R 4.3.2 for all analyses. Generalized linear mixed models (GLMMs) were used to analyse the relationship between *S. anglicus* detections and treatment years for each of the 13 search areas. ‘Year’ was included as a continuous fixed effect and ‘search area’ as a categorical fixed effect. ‘Search block’ was included as a random effect to account for the non-independence of observations made within the same search block over different years. Year was transformed by subtracting 2000 to improve model convergence by reducing the numerical range. *S. anglicus* detections (with pseudo-zero values) were used as the response variable, with a binomial distribution in a GLMM using the R package ‘lme4’ (Bates, 2015). A second model was run using the ‘true’ data (sans zero-values) to assess the effect of less regular annual search efforts on future detection probabilities (Table S2, complete data tables). However, our results focus on the pseudo-zero model and its more realistic outputs.

We used delta marginal R-squared (R^2^m) to assess model fit, with a resulting value of 0.26, indicating that over a quarter of the variation in the data could be explained by the model’s fixed effects. While modest, this is typical of ecological field data and still captures key drivers. Model predictions were generated from 2013 to 2040, with confidence intervals set to 99%, using the ‘ggeffects’ package in R (Lüdecke, 2018). We analysed our results using the model predictions’ summary tables, to two decimal places. The year of functional eradication for each search area was determined as the first year that the likelihood of detecting *S. anglicus* is predicted to be <0.01, with the 99% upper confidence limit being <0.01. Conservative confidence intervals were necessary in the absence of a plant detectability estimate.

### Model validation and sensitivity testing

We used two methods to cross-validate our results. First, we tested the robustness of the model’s predictions by subsetting the data to 2020 and generating predictions to 2024. We compared these predictions to the observed data, to determine if the observed data fell within the subsetted model’s confidence intervals.

Second, we tested model sensitivity, addressing the model’s assumption of zero future detections. We focussed sensitivity analyses on the ‘Kaituna’ search area, which recorded the highest likelihood of future detections, providing a ‘worst case scenario’ for testing. We simulated detections beyond the dataset, from 2025 to 2030, to evaluate how this impacted model predictions. We introduced both single annual detections and compounded detections in consecutive years, to assess potential delays in the predicted functional eradication year.

Simulated detections included up to five consecutive finds in the ‘Kaituna’ search block through to 2030. Detections exceeding this number would strongly suggest a potential source population outside the defined search blocks, making it unnecessary to extend the model beyond these limits, as such data would no longer pertain to an eradication framework. Moreover, testing solely within ‘Kaituna’ highlights a core limitation of the model: the interconnectivity of marine areas prevents complete independence between blocks, despite the inclusion of search blocks as a random effect. Although this limitation is unavoidable, our use of the 13 larger search areas for analysis counters some of the non-independence, while providing a useful way to interpret the interconnected waterway in terms of *S. anglicus* spread to and from adjacent areas.

## Results

Since 2013, there were 162 *S. anglicus* detections, with over half occurring in the first three years of the dataset. Of the 907 total observations (including pseudo-zero datapoints), 109 showed detection, predominantly as single-plant detections within a search block for a given year. Had every search block been surveyed annually, the dataset would have included 948 datapoints. Between 2019 and 2024—representing the most recent half of the dataset—only 17 plants (15.6% of all detections) were found across all search areas.

Our model demonstrated that annual control efforts have a significant effect on *S. anglicus* detections (P <0.0001). Model results showed a negative relationship between year and detections, with coefficients across models ranging from -0.39 to -0.45 log-odds ratio (Table 1). Thus, detections decreased with each control season since 2013.

**Table 1:** Model coefficients, with standard errors (SEM, n = 907 replicates for the pseudo-zero dataset, n = 500 for the ‘true’/observed dataset) and P-values for Spartina detections across years.

| Model | Coefficient | SEM | P-value |
| --- | --- | --- | --- |
| 1 \| Presence/absence, pseudo-zeros data | -0.45 (log-odds ratio) | 0.052 | <0.0001 |
| 2 \| Presence/absence, true data | -0.39 (log-odds ratio) | 0.054 | <0.0001 |

Model predictions indicated that *S. anglicus* detections will regress to functional eradication under the current management regime, with the pseudo-zero model predicting eradication by 2032, at an upper confidence limit of <0.01 (Table 2). This projection is 7 years from the current year (2025) and over half the length of the total dataset, underscoring the long confirmation phase characteristic of near eradication. The ‘true’ dataset (without pseudo-zero values) produced more distant predictions, estimating eradication by 2036 (Table 2, ‘Dataset 2’). This result reflects the smaller number of observations (n=500) with a higher proportion of detections.

**Table 2:** GLMM model predictions of eradication year for each search area. Eradication years were determined by the first year with a 99% upper confidence limit of <0.01 log-odds ratio. Highlighted in yellow is the most conservative year of predicted eradication, determined by the most persistent search area, ‘Kaituna’. Marginal R-squared values (R^2^m) are reported to assess model fit. Delta R^2^m was chosen for its balanced evaluation of model fit.

| Area | Dataset 1: zero/PA | Dataset 2: true/PA |
| --- | --- | --- |
| Pelorus back | 2025 | 2032 |
| Pelorus central | 2029 | 2033 |
| Pelorus | 2031 | 2034 |
| Brownlees | 2028 | 2033 |
| Kaikumera | 2029 | 2033 |
| Kaituna | 2032 | 2036 |
| Mahakipawa | 2029 | 2033 |
| Kaiuma | 2032 | 2035 |
| Mahau | 2031 | 2035 |
| Kenepuru | 2026 | 3031 |
| Inner sounds | 2027 | 3031 |
| Outer sounds | 2026 | 2034 |
| Tennyson | 2026 | 2030 |
| **Delta $R^2_m$** | 0.26 | 0.26 |

### Model Testing

Cross-validation using an earlier subset of data to predict the likelihood of detection for the years 2020 to 2024 demonstrates that the projected regression pattern aligns well with the observed data, although with several values falling outside the confidence intervals (Figure 4). Sensitivity testing showed that compound future detections in the ‘Kaituna’ area would delay functional eradication incrementally with each additional simulated detection. Specifically, five detections per year from 2025 to 2030 resulted in a functional eradication year of 2037, five years later than our primary model’s projected year of 2032 (Table 3). In contrast, a single simulated detection in ‘Kaituna’ in any given year between 2025 and 2030 increased the predicted functional eradication date by just one year, remaining constant from 2033 thereafter.

**Figure 4:**
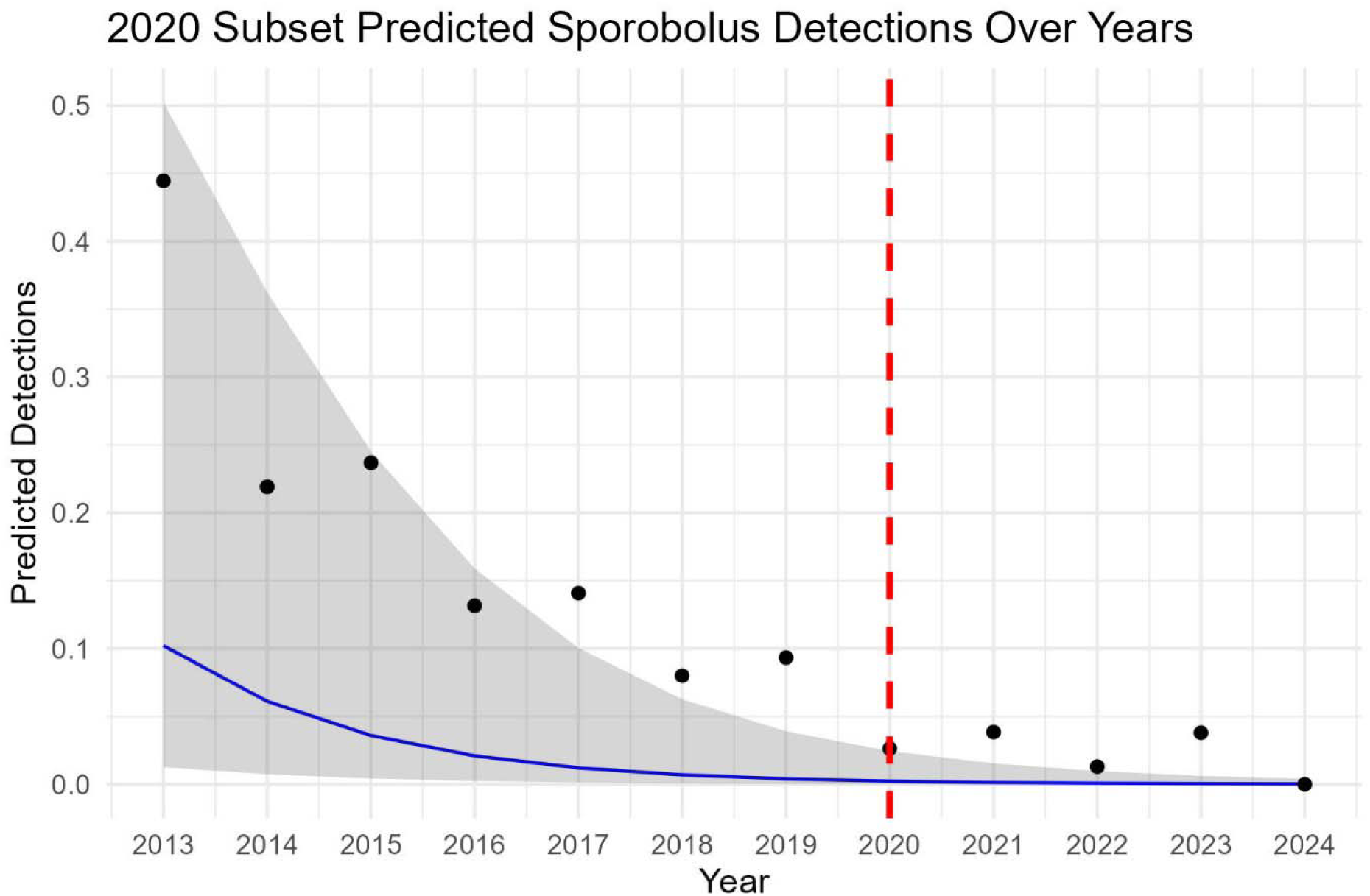
Subsetted cordgrass dataset from 2013 to 2020, and the model’s predicted values from 2020 to 2024, with 99% confidence intervals shown as a shaded region. Observed datapoints from 2013 to 2024 are displayed as dots, while the red dashed line marks the 2020 subset boundary. The y-axis is the predicted likelihood of detection in a given year.

**Table 3:** Model predictions for simulated detections in consecutive years, compared to our primary model results (‘original data’, predicted eradication 2032). All simulations represent single detections per listed year/s, e.g. ‘2025 - 2027’ represents a single simulated detection in 2025, 2026 and 2027, with the model forecasting a more remote eradication year of 2035 with these additional detections.

| Year/s of simulated detections | Predicted year of eradication |
| --- | --- |
| Original data | 2032 |
| 2025 | 2033 |
| 2025 - 2026 | 2034 |
| 2025 - 2027 | 2035 |
| 2025 - 2028 | 2035 |
| 2025 - 2029 | 2036 |
| 2025 - 2030 | 2037 |

## Discussion

The results of our modelling provide insight into how the detection probability of *Sporobolus anglicus* has changed over time in response to annual control in Te Hoiere, Marlborough Sounds, directly addressing our primary aim. Model predictions support the evaluation of long-term management outcomes, by painting a picture of the project’s trajectory toward functional eradication, thus meeting our broader objectives.

The model clearly demonstrates that current management strategies are working, with the likelihood of future detections falling below 0.01 in all search areas by 2032 (Figure S2). The significant negative response of detections across years, coupled with the absence of new detections in 2024, suggests any remaining plants are either sterile or present in very low numbers. This aligns with known traits of *S. anglicus,* which is highly fecund and liable to tidal spread (Marks & Truscott, 1985). If seed events were occurring, we would expect to detect more plants in the surveyed areas, especially given the regularity of surveillance.

The ‘Kaituna’ search area was the most persistent for *S. anglicus*, reaching our defined threshold for declaring functional eradication by 2032. This sets a conservative benchmark for other search areas. By also modelling the ‘true’ dataset (excluding pseudo-zero values), we provide an additional appraisal of the project’s trajectory, in view of practical survey limitations. This model projected a delayed functional eradication timeline, extending to 2036 (Figure S2). The higher number of zeros in the pseudo-zero dataset leads the model to more readily predict future *S. anglicus* non-detections. While this approach may yield more realistic results, we cannot guarantee true absence in cases where values were substituted; we have instead made informed assumptions of non-detections based on previous detections.

To account for this uncertainty, we recommend that decision-making takes into consideration projections from both models. Importantly, the four-year difference between them strongly underscores the need for intensified surveillance in the coming years. Confirming non-detections across all search areas will be key to confidently retire the project with statistical support.

This is exemplified in the ‘Pelorus’ search area, which yielded precise predictions of detection probability, suggesting functional eradication by 2031. ‘Pelorus’ has been regularly searched since 2013, with few missing values and most records showing either absence or isolated detections within a given season. In contrast, the ‘Brownlees’ and ‘Outer Sounds’ areas showed the greatest disparity between the pseudo-zero and ‘true’ models, likely because detections were concentrated in a single block within these areas (Table S2, complete data tables). For example, detections in ‘Outer Sounds’ were limited to the ‘Manaroa’ block, while the six remaining blocks yielded no detections across two surveys each since 2017. Despite the number of missing data in other years, repeated non-detections in real surveys validates our use of substituted zero-values, lending confidence to the accuracy of the pseudo-zero model’s predictions.

According to the pseudo-zero model, functional eradication in ‘Outer Sounds’ could occur as early as 2026, whereas the ‘true’ dataset projects 2034 – an 8-year difference. These divergent timelines offer practical insights in allocating field resources. Given the limited and localised detections, ‘Outer Sounds’ may be ready to transition into a monitoring phase with more targeted surveillance, allowing search efforts to focus on more persistent areas such as ‘Kaituna’ and ‘Pelorus Central’.

However, model results must still be interpreted within an ecological context (Panetta, 2015), especially given the connectedness of the Te Hoiere waterways. While tidal dynamics were not explicitly modelled, including search blocks as a random effect helped account for spatial proximity within and between grouped blocks. With seed-holding periods ranging from 6 months to 4 years for *Sporobolus* species (Marks & Truscott, 1985; Infante-Izquierdo et al., 2023; Hubbard 1970; Xiao et al., 2010), project withdrawal should only follow four consecutive years of recorded absence across all potential habitat (Panetta, 2007). While our projections meet this threshold, increasing search frequency would strengthen confidence by confirming non-detections and reducing the chance of missed plants. We recognise that *S. anglicus* may take one or more years to be visible to searchers, due to inconspicuous or young plants, or searcher distraction. To account for imperfect detectability, we applied conservative confidence intervals as a buffer against false negatives.

Despite the risk of detection bias and the lack of a detectability estimate in our models, several features of the target plant and the method of survey in the Te Hoiere project suggest that, through a practical lens, the risk of a missed plant may be lower than in other contexts. The close-set survey method (Figure 3) typically places three searchers within view of a single survey line, increasing the chance of detection. This is further enhanced by the slow pace of survey which allows greater visual scrutiny (C. Wadsworth, pers. comm., May 2024). Additionally, detectability improves under certain field conditions, particularly when the target species contrasts markedly with surrounding vegetation, when floral diversity is low, and when vegetation structure is uniform (Hauser et al. 2022). In Te Hoiere, *S. anglicus* has typically appeared in unvarying hip-height reedbeds dominated by *Apodasmia similis* and *Juncus krausiana* - both visually distinct in colour and form from *S. anglicus* (personal experience). These factors are likely to improve the reliability of visual surveys in this setting.

### Model Assumptions

The predictive model primarily assumes that no additional plants will be found. While the raw data and model regression strongly support this, results should be interpreted with caution due to limitations of the current search program, which does not fully meet Panetta’s laws of weed eradication (Panetta & Brooks, 2008). Namely, (A) not all potential habitats are consistently surveyed, (B) the target area may not be isolated from external weed ingress, and (C) detection is less than near-certain. While comprehensive, the Te Hoiere eradication program does not include all potential *S. anglicus* habitat. Although delimitations of coastlines and reedbeds without a history of *S. anglicus* have been intermittently undertaken, single records of non-detection do not preclude later establishment events from dormant seed or known infestations. Furthermore, there remains a minor risk of *S. anglicus* being inadvertently or intentionally introduced to the waterway.

Additionally, the model assumes that search effort and plant detectability is constant across years. While total surveillance time varied little between seasons, search teams have different personnel each year, which may impact detection rates (Ng & Driscoll, 2015; Couvreur et al., 2015). However, the presence of one or more long-term team members ensures some consistency (Bornand, 2014). Crucially, the Field Team Lead has been consistent for the duration of the dataset.

Despite our use of conservative model parameters, ecological forecasting remains inherently uncertain (Oliver & Roy, 2015). We used linear regression for *S. anglicus* detections, aligning with documented weed reductions from consistent removal efforts (Panetta et al., 2011). Our model therefore forecasts expected detections, assuming current search parameters remain unchanged. However, it does not directly predict presence or absence. While indirect, this serves as a reliable proxy for functional eradication, by quantifying the usefulness of continuing with manual surveillance under the current regime, when further detections are extremely unlikely.

Subset and sensitivity analyses showed the model to be stable. While subset analysis did not fully capture the observed values up to 2024, the model demonstrated consistent patterns of decline between the original and subsetted datasets, aligning well with the known trend. Sensitivity analyses indicated that a single positive detection in the next five years would minimally impact the eradication timeline. A solitary detection in future years is likely to be outlier and may not significantly impact the overall downward trend toward consistent records of absence predicted by the model. This is biologically reasonable; for instance, a single detection in 2030 following five consecutive years without detections would indicate a regrowth event or an improbable seed-holding scenario, rather than a persistent source infestation capable of altering the project’s trajectory and invalidating the model’s assumptions. Conversely, consecutive detections would significantly prolong the program, with the model extending the tail end of projected eradication. Such a situation could indicate a persistent source population, and should trigger a review of the project’s aims and strategies.

### Management Recommendations

Based on our analyses, we recommend several management strategies to help achieve functional *S. anglicus* eradication in the Marlborough Sounds within the projected timeline of 2032. We strongly suggest that these strategies be applied in other *Sporobolus* eradication programs in Aotearoa and abroad, especially those approaching low-density stages.

*1) Increased search frequency*. We recommend annual or biennial searches of all blocks for a five-year intensive monitoring period at the tail-end of eradication efforts. This schedule addresses the 2–3-year lag between germination and seed set (Gray & Benham, 1990) and aligns with known seed-holding durations (Marks & Truscott, 1985; Strong & Ayres, 2013). The rapid growth and high fertility of *S. anglicus* make infrequent searches inadequate, and under the current regime in Te Hoiere, some blocks go unsearched for up to three years. This delay increases the risk of seed set and may prolong the plateau phase (Panetta & Lawes, 2005). Short-term cost increases could be offset by avoiding the costs of drawn-out management (Panetta, 1999). More regular surveys also reduce the influence of environmental factors on detectability.
*2) Expanded delimitation*. All potential habitats should be surveyed at least twice within this five-year period, including shorelines, reedbeds, and riverbanks up to 2km upstream of river mouths, where salinity remains suitable for *S. anglicus* persistence (Brown & Raal, 2013; Partridge, 1987). Complete coverage is essential (Panetta, 2007). In the dense reedbeds typical of the upper estuarine zone, drones (Esposito et al., 2021) and *Sporobolus*-trained detection dogs (Goodwin et al., 2010) can supplement grid-search efforts; both have shown promise in detecting small or cryptic individuals (Brown & Raal, 2013).
*3) Staff consistency*. Successful weed projects rely on experienced, dedicated personnel who advocate for the program, secure funding, and perform reliable fieldwork (Brown & Raal, 2013; Howell, 2012; Bornand, 2014). Familiarity with search blocks also reduces training time and improves data consistency. In Te Hoiere, several field staff have participated for multiple seasons, bringing local knowledge and stalwart commitment to the goal of eradication.
*4) Adaptive management.* Model predictions should be updated biannually, refining projections and accounting for potential new source populations. More consistent surveillance will improve model accuracy. A significant discovery outside current search areas, such as a seeding plant, should prompt a reassessment of strategies, particularly given the need for high confidence that no sexual or vegetative reproduction is occurring within the catchment (Panetta & Lawes, 2005).
*5) Environmental DNA.* Environmental DNA (eDNA) could offer a sensitive and cost-effective tool for post-eradication monitoring (Lodge et al., 2012). By sampling waterways and comparing results to reference libraries, qPCR could be used to confirm absence (Gantz et al., 2018; Anglès d’Auriac et al., 2019; Zhu et al., 2024). This could serve as a valuable backstop, addressing concerns around premature project retirement. However, legacy DNA from rhizomatous material may confound results (Zhu et al., 2024; Collins et al., 2018). Targeted research is needed across sites with varying *Sporobolus* densities, to develop a species-specific protocol suited to both low-density and post-eradication stages, enhancing long-term monitoring capacity.
*6) Detectability estimates*. Estimating detectability using re-survey trials was not feasible in this study - a limitation common to many weed control programs (e.g. West & Havell, 2019; Brown & Brown, 2015). However, future research conducted independently of operational constraints could enable detectability trials in representative reedbeds. Using a standardised grid-search method, such partnerships could yield species-specific estimates and improve model accuracy.

## Conclusion

Under the current management regime, our modelling predicts that *S. anglicus* detections in Te Hoiere should decline to levels consistent with functional eradication by 2032. Drawing on real-world detection data, this approach offers a pragmatic forecasting tool to support decision-making. It highlights when continued surveillance is unlikely to yield further detections, and provides detection probabilities for 13 geographic zones, helping managers to prioritise searches as project retirement appears on the horizon.

To maintain progress toward eradication, we advocate for full delimitation of all suitable habitat within Te Hoiere, to ensure no residual plants fall outside active management. If spread has been effectively halted, and each search block is revisited annually or biennially, eradication within the projected timeframe remains achievable.

Regardless of increased efforts, however, perfect detection can never be guaranteed. As Rout (2017) notes, managers must ultimately “sit down and decide what is acceptable” using the tools available. While our results support a confident approach toward functional eradication, absolute certainty of absence is unattainable. Decisions must therefore balance technical feasibility with practical risk tolerance.

Nonetheless, this case study offers a replicable framework for other *Sporobolus* eradication initiatives in Aotearoa. Our results underscore the need for sustained effort in the final stages of surveillance, in aid of a confident exit from what is an undoubtedly successful control program. Weed eradication requires constant pressure on the accelerator, and a final decisive push to ensure the goal is met. The next five years will be critical for either accelerating or prolonging *S. anglicus* eradication efforts in the Marlborough Sounds. If our recommendations are implemented, it is likely that functional eradication should occur within our projected timeline.

## Supporting information

Supplemental Table 2: Complete Data Tables

## Acknowledgments

We acknowledge Ngāti Kuia, mana whenua and kaitiaki of Te Hoiere. *N*ā*u te rourou*.

Additionally, we thank the Department of Conservation (DOC) Sounds Area Office for providing data. Thank you to DOC rangers Daniel Palmer, Philip Clerke and Clinton Wadsworth for providing personal observations. Thank you also to DOC weed advisor Kate McAlpine, who we consulted on *Sporobolus* taxonomy.

A special mention is needed for a few long-serving “Spartinians” with whom the primary author worked shoulder-to-shoulder with in the foetid mudflats of Te Hoiere in 2017: Gary (Gazza Gazelle), Marc (Marco Polo), Jacko (just Jacko) and Clinton (Crimson Wisdom). A passion for weed eradication was born in that fateful season. May the reeds long prick your knees. *Ng*ā *mihi nunui ki a koutou, ng*ā*i patu t*ō*t*ō.

## Declarations

The authors received no funding for this research and declare no conflicts of interest.

## Data and Code Availability

The data used in this study are available upon request from the NZ Department of Conservation (DOC), Marlborough Sounds Area Office.

Code is available upon request from the primary author.

## Appendix

**Table S1:** Summary of annual spartina detections and searched blocks.

| Year | Total No. Plants/Sites | No. Blocks Searched |
| --- | --- | --- |
| 2009 | 507 | NA |
| 2010 | 417 | NA |
| 2011 | 114 | NA |
| 2012 | 66 | NA |
| 2013 | 59 | 48 |
| 2014 | 20 | 45 |
| 2015 | 22 | 52 |
| 2016 | 16 | 50 |
| 2017 | 15 | 59 |
| 2018 | 14 | 54 |
| 2019 | 8 | 28 |
| 2020 | 2 | 44 |
| 2021 | 3 | 45 |
| 2022 | 1 | 31 |
| 2023 | 3 | 39 |
| 2024 | 0 | 28 |

**Figure S1:**
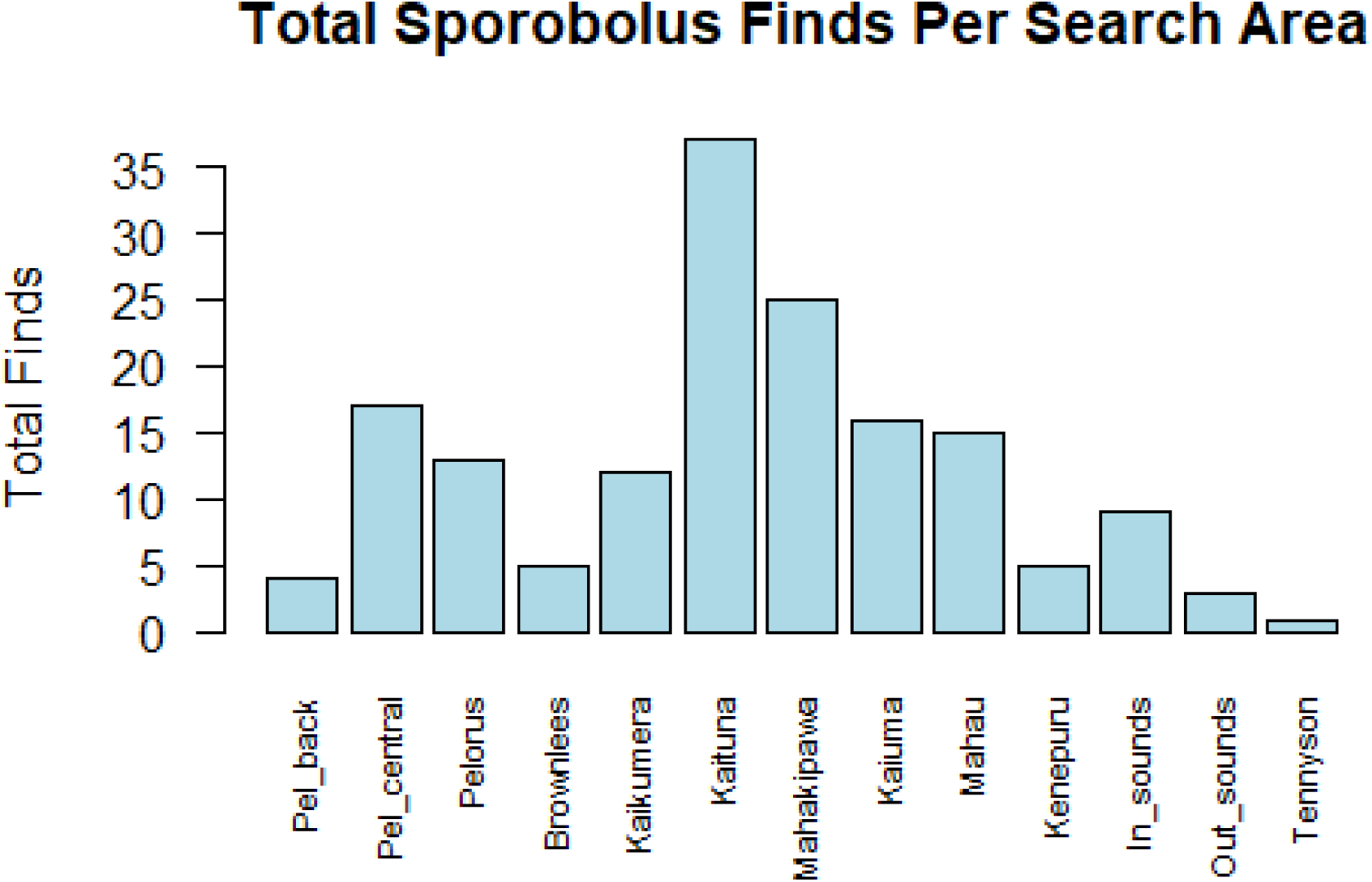
Total *S. anglicus* detections per search area from 2013 to 2024. Search areas are listed from the furthest upstream (‘Pelorus back’, far left of graph) to the furthest downstream (‘Tennyson’, far right of graph).‘Kaituna’, with the most detections, may be having downstream impacts on the ‘Mahakipawa’, ‘Kaiuma’ and ‘Mahau’ areas.

**Figure S2:**
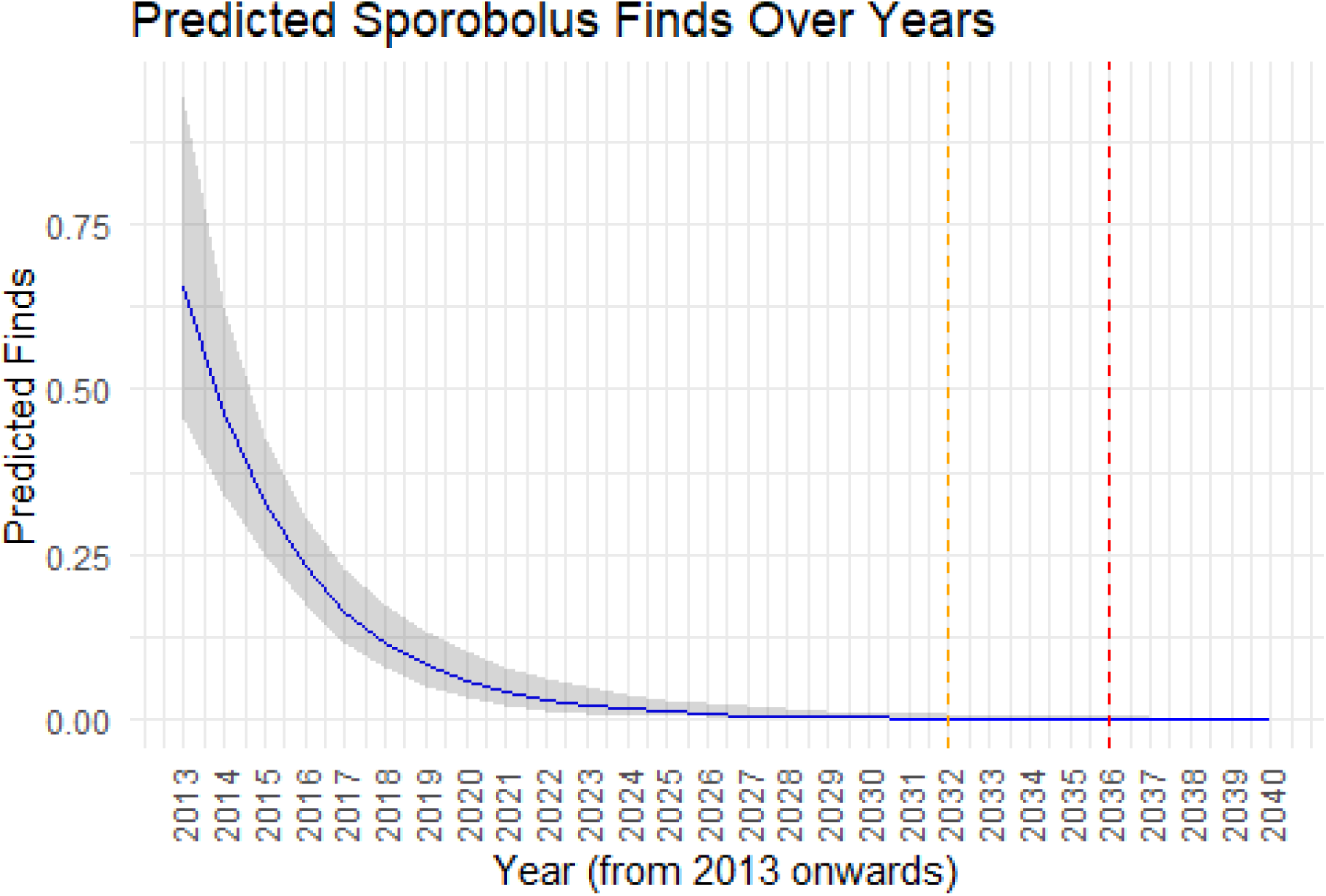
GLMM-predicted *S. anglicus* detections with all search areas aggregated. The 99% confidence intervals are shown by the grey shaded area. The orange dashed line represents the predicted eradication year of 2032 modelled with pseudo-zeros data. The red dashed line represents the conservative predicted eradication year of 2036 modelled with the observed data.

*[Appendix: Table S2 in separate document due to large size and alternative formatting]*

## Resources

Ainouche, M., & Gray, A. (2016). Invasive Spartina: lessons and challenges. Biological Invasions, 18, 2119–2122.

Anglès d’Auriac, M. B., Strand, D. A., Mjelde, M., Demars, B. O., & Thaulow, J. (2019). Detection of an invasive aquatic plant in natural water bodies using environmental DNA. PloS one, 14(7), e0219700.

Bates, D. M., Martin; Bolker, Ben; Walker, Steve. (2015). Fitting Linear Mixed-Effects Models Using lme4. Journal of Statistical Software, 67. doi:10.18637/jss.v067.i01.

Bowie, I. J. S. (1963). Land utilisation in the Marlborough Sounds.

Bornand, C. N., Kéry, M., Bueche, L., & Fischer, M. (2014). Hide□and□seek in vegetation: time□to□detection is an efficient design for estimating detectability and occurrence. Methods in Ecology and Evolution, 5(5), 433–442.

Brown, K., & Brown, D. (2015). Control to eradication of Tradescantia fluminensis on Stephens Island (Takapourewa): the importance of systematic and persistent effort. DOC Research & Development Series, 15.

Brown, K., & Raal, P. (2013). Is eradication of spartina from the South Island feasible*?* Department of Conservation.

Collins, R. A., Wangensteen, O. S., O’Gorman, E. J., Mariani, S., Sims, D. W., & Genner, M. J. (2018). Persistence of environmental DNA in marine systems. Communications Biology, 1(1), 185.

Marlborough District Council. (2024). Regional Pest Management Plan. https://www.marlborough.govt.nz/repository/libraries/id:2ifzri1o01cxbymxkvwz/hierarchy/documents/environment/regional-pest-management-plan/Regional_Pest_Management_Plan

Couvreur, J. M., Fiévet, V., Smits, Q., & Dufrêne, M. (2015). Evaluation of observer effect in botanical surveys of grasslands. *Biotechnologie, Agronomie*, Société et Environnement.

Department of Conservation. (2016). War on weeds. https://www.beehive.govt.nz/sites/default/files/War-on-Weeds.pdf

Esposito, M., Crimaldi, M., Cirillo, V., Sarghini, F., & Maggio, A. (2021). Drone and sensor technology for sustainable weed management: A review. Chemical and Biological Technologies in Agriculture, 8, 1–11.

Gantz, C. A., Renshaw, M. A., Erickson, D., Lodge, D. M., & Egan, S. P. (2018). Environmental DNA detection of aquatic invasive plants in lab mesocosm and natural field conditions. Biological Invasions, 20, 2535–2552.

Goodwin, K. M., Engel, R. E., & Weaver, D. K. (2010). Trained dogs outperform human surveyors in the detection of rare spotted knapweed (Centaurea stoebe). Invasive Plant Science and Management, 3(2), 113–121.

Gray, A. J., & Benham, P. E. (1990). Spartina anglica-a research review. HMSO.

Green, S. J., & Grosholz, E. D. (2021). Functional eradication as a framework for invasive species control. Frontiers in Ecology and the Environment, 19(2), 98–107.

Hammond, M., & Cooper, A. (2002). Spartina anglica eradication and inter-tidal recovery in Northern Ireland estuaries. Turning the tide: the eradication of invasive species, 124–131.

Harrison, J. B., Sunday, J. M., & Rogers, S. M. (2019). Predicting the fate of eDNA in the environment and implications for studying biodiversity. Proceedings of the Royal Society B, 286(1915), 20191409.

Harrison Day, V., Prahalad, V., McHenry, M. T., Aalders, J., & Kirkpatrick, J. B. (2023). Introduced Spartina anglica modifies fish habitat in southern temperate succulent saltmarshes. Restoration Ecology, 31(7), e13812.

Howell, C. J. (2012). Progress toward environmental weed eradication in New Zealand. Invasive Plant Science and Management, 5(2), 249–258.

Hubbard, J. (1970). Effects of cutting and seed production in Spartina anglica. Journal of Ecology, 58(2), 329–334.

Hulme, P. E. (2012). Weed risk assessment: a way forward or a waste of time?. Journal of Applied Ecology, 49(1), 10–19.

Hulme, P. E. (2020). Plant invasions in New Zealand: global lessons in prevention, eradication and control. Biological invasions, 22(5), 1539–1562.

Hume, T., Gerbeaux, P., Hart, D., Kettles, H., & Neale, D. (2016). A classification of New Zealand’s coastal hydrosystems.

Infante-Izquierdo, M. D., Romero-Martín, R., Castillo, J. M., Grewell, B. J., Soriano, J. J., Nieva, F. J. J., & Muñoz-Rodríguez, A. F. (2023). Seed viability, spikelet dispersal, seed banks and seed storage requirements for native and invasive cordgrasses (Genus spartina) in Southwest Iberian Peninsula. Wetlands, 43(1), 8.

Kerr, D. W., Hogle, I. B., Ort, B. S., & Thornton, W. J. (2016). A review of 15 years of Spartina management in the San Francisco Estuary. Biological invasions, 18(8), 2247–2266.

Kriwoken, L. K., & Hedge, P. (2000). Exotic species and estuaries: managing Spartina anglica in Tasmania, Australia. Ocean & Coastal Management, 43(7), 573–584.

Lee, W. G., & Partridge, T. R. (1983). Rates of spread of Spartina anglica and sediment accretion in the New River Estuary, Invercargill, New Zealand. New Zealand Journal of Botany, 21(3), 231–236.

Lodge, D. M., Turner, C. R., Jerde, C. L., Barnes, M. A., Chadderton, L., Egan, S. P., … & Pfrender, M. E. (2012). Conservation in a cup of water: estimating biodiversity and population abundance from environmental DNA. Molecular ecology, 21(11), 2555–2558.

Loebl, M., van Beusekom, J. E., & Reise, K. (2006). Is spread of the neophyte Spartina anglica recently enhanced by increasing temperatures? Aquatic Ecology, 40(3), 315–324.

Lu, K., Han, G., & Wu, H. (2022). Effects of Spartina alterniflora invasion on the benthic invertebrate community in intertidal wetlands. Ecosphere, 13(3), e3963.

Lüdecke, D. (2018). ggeffects: Tidy Data Frames of Marginal Effects from Regression Models. Journal of Open Source Software, 3(26). 10.21105/joss.00772

Lyu, C., Zhang, S., Ren, X., Liu, M., Leung, K. S. K., He, T., … & Choi, C. Y. (2023). The effect of Spartina alterniflora eradication on waterbirds and benthic organisms. Restoration Ecology, 31(8), e14023.

MacKenzie, D. I., Nichols, J. D., Lachman, G. B., Droege, S., Andrew Royle, J., & Langtimm, C. A. (2002). Estimating site occupancy rates when detection probabilities are less than one. Ecology, 83(8), 2248–2255.

Marks, T., & Truscott, A. (1985). Variation in seed production and germination of Spartina anglica within a zoned saltmarsh. The Journal of Ecology, 695–705.

Marlborough District Council (2024). Regional Pest Management Plan. https://www.marlborough.govt.nz/repository/libraries/id:2ifzri1o01cxbymxkvwz/hierarchy/documents/environment/regional-pest-management-plan/Regional_Pest_Management_Plan

McAlpine, K. G., & Howell, C. J. (2024). List of environmental weeds in New Zealand 2024. Science for Conservation, 340.

Millen, P., & Waddel, S. (2003). Resource Consent Application for the Control of Spartina Grass in Havelock Estuary.

Milne, D. H. (2007). Controlling an invasive salt marsh grass (Spartina patens) in Washington State: a case study of resilience. Environmental Practice, 9(4), 251–265.

Moore, J. L., Hauser, C. E., Bear, J. L., Williams, N. S., & McCarthy, M. A. (2011). Estimating detection–effort curves for plants using search experiments. Ecological Applications, 21(2), 601–607.

Muhammad, B. L., & Ki, J. S. (2024). Polyploidy and genome evolution in the common cordgrass Spartina anglica: an enigmatic evolution of allopolyploidy. Systematics and Biodiversity, 22(1), 2357091.

Mullins, P., & Marks, T. (1987). Flowering phenology and seed production of Spartina anglica. The Journal of Ecology, 1037–1048.

Ng, K., & Driscoll, D. A. (2015). Detectability of the global weed Hypochaeris radicata is influenced by species, environment and observer characteristics. Journal of Plant Ecology, 8(4), 449–455.

Oliver, T. H., & Roy, D. B. (2015). The pitfalls of ecological forecasting. Biological Journal of the Linnean Society, 115(3), 767–778. 10.1111/bij.12579

Panetta, F. (1999). Can we afford to delay action against weeds in valued natural areas?

Panetta, F. (2015). Weed eradication feasibility: lessons of the 21st century. Weed Research, 55(3), 226–238.

Panetta, F. D. (2007). Evaluation of weed eradication programs: containment and extirpation. Diversity and Distributions, 13(1), 33–41.

Panetta, F. D. and S. J. Brooks (2008). Evaluating progress in weed eradication programs. Pages 418–420 in Proceedings of the 16th Australian Weeds Conference. Brisbane: Queensland Weeds Society.

Panetta, F. D., Cacho, O., Hester, S., Sims Chilton, N., & Brooks, S. (2011). Estimating and influencing the duration of weed eradication programmes. Journal of Applied Ecology, 48(4), 980–988.

Panetta, F. D., & Lawes, R. (2005). Evaluation of weed eradication programs: the delimitation of extent. Diversity and Distributions, 11(5), 435–442.

Partridge, T. (1987). Spartina in New Zealand. New Zealand Journal of Botany, 25(4), 567–575.

Patten, K., O’Casey, C., & Metzger, C. (2017). Large-scale chemical control of smooth cordgrass (Spartina alterniflora) in Willapa Bay, WA: towards eradication and ecological restoration. Invasive Plant Science and Management, 10(3), 284–292.

Pyšek, P., & Richardson, D. M. (2010). Invasive species, environmental change and management, and health. Annual review of environment and resources, 35, 25–55.

Ramsey, D. S., Anderson, D. P., & Gormley, A. M. (2023). Invasive species eradication: How do we declare success?. Cambridge Prisms: Extinction, 1, e4.

Reeder, T. G., & Hacker, S. D. (2004). Factors contributing to the removal of a marine grass invader (Spartina anglica) and subsequent potential for habitat restoration. Estuaries, 27, 244–252.

Regan, T. J., McCarthy, M. A., Baxter, P. W., Dane Panetta, F., & Possingham, H. P. (2006). Optimal eradication: when to stop looking for an invasive plant. Ecology letters, 9(7), 759–766.

Rejmánek, M., & Pitcairn, M. J. (2002). When is eradication of exotic pest plants a realistic goal. Turning the tide: the eradication of invasive species, 249–253.

Rout, T. M. (2017). Declaring Eradication of an Invasive Species. In A. P. Robinson, T. Walshe, M. A. Burgman, & M. Nunn (Eds.), Invasive Species: Risk Assessment and Management (pp. 334–347). chapter, Cambridge: Cambridge University Press.

Sheehan, M. R. (2008). Assessment of the Potential Impacts of Large-scale Eradication of Spartina anglica from the Tamar Estuary (Doctoral dissertation, University of Tasmania).

Sheehan, M. R., & Ellison, J. C. (2014). Intertidal morphology change following Spartina anglica introduction, Tamar Estuary, Tasmania. Estuarine, Coastal and Shelf Science, 149, 24–37.

Sheehan, M. R., & Ellison, J. C. (2015). Tidal marsh erosion and accretion trends following invasive species removal, Tamar Estuary, Tasmania. Estuarine, Coastal and Shelf Science, 164, 46–55.

Shimeta, J., Saint, L., Verspaandonk, E. R., Nugegoda, D., & Howe, S. (2016). Long-term ecological consequences of herbicide treatment to control the invasive grass, Spartina anglica, in an Australian saltmarsh. Estuarine, Coastal and Shelf Science, 176, 58–66.

Sileshi, G., Hailu, G., & Nyadzi, G. I. (2009). Traditional occupancy–abundance models are inadequate for zero-inflated ecological count data. Ecological Modelling, 220(15), 1764–1775. 10.1016/j.ecolmodel.2009.03.024.

Simberloff, D. (2003). Eradication—preventing invasions at the outset. Weed Science, 51(2), 247–253.

Strong, D. R., & Ayres, D. R. (2013). Ecological and evolutionary misadventures of Spartina. Annual review of ecology, evolution, and systematics, 44, 389–410.

Tyre, A. J., Tenhumberg, B., Field, S. A., Niejalke, D., Parris, K., & Possingham, H. P. (2003). Improving precision and reducing bias in biological surveys: estimating false negative error rates. Ecological Applications, 13(6), 1790–1801.

Underwood, J. (2021). Marlborough District Council Biosecurity Operational Plan 2018-2028 (amended 16 Sept 2021).

Urlich, S. C., & Handley, S. J. (2020). From ‘clean and green’to ‘brown and down’: A synthesis of historical changes to biodiversity and marine ecosystems in the Marlborough Sounds, New Zealand. Ocean & Coastal Management, 198, 105349.

Wang, S., Martin, P. A., Hao, Y., Sutherland, W. J., Shackelford, G. E., Wu, J., … & Li, B. (2023). A global synthesis of the effectiveness and ecological impacts of management interventions for Spartina species. Frontiers of Environmental Science & Engineering, 17(11), 141.

West, C. J., & Havell, D. (2019). Weed eradication on Raoul Island, Kermadec Islands, New Zealand: progress and prognosis. Island invasives: scaling up to meet the challenge, (62), 435.

Wintle, B., Garrard, G., & Bekessy, S. (2009). Determining necessary survey effort to detect invasive weeds in native vegetation communities. https://cebra.unimelb.edu.au/data/assets/pdf_file/0009/2220786/0906-final-report.pdf

Xiao, Y., Tang, J., Qing, H., Ouyang, Y., Zhao, Y., Zhou, C., & An, S. (2010). Clonal integration enhances flood tolerance of Spartina alterniflora daughter ramets. Aquatic Botany, 92(1), 9–13.

Xu, D. F., Yuan, Q., Lu, L. W., Tan, B., Ge, M., Chen, J. Y., … & Zhao, B. (2024). Is Spartina alterniflora eradication project in Chongming Island a nature-based solution?. Nature-Based Solutions, 6, 100178.

Zhu, X., Bell, K. L., Wu, H., & Gopurenko, D. (2024). Development of an Environmental DNA Assay for Prohibited Matter Weed Amazon Frogbit (Limnobium laevigatum). Environments, 11(4), 66.

