## Supplemental Table 2: Complete Data Tables for "Modelling the decline in *Sporobolus anglicus* detections toward functional eradication: a case study in the Marlborough Sounds, New Zealand"

*Appendix*

**Table S2:** Complete data tables of GLMM results with predicted *S. anglicus* detections from 2013 until 2040, including 99% confidence intervals (CI) and standard errors (SEM in all cases; n = 907 replicates for the pseudo-zero data, n = 500 replicates for the observed data).

| | **True data, count data** | | | | | **True data, presence/absence data** | | | | | **Zero data, count data** | | | | | **Zero data, presence/absence data** | | | | |
|---|---|---|---|---|---|---|---|---|---|---|---|---|---|---|---|---|---|---|---|---|
| **Group:** | Year | pred. | CI.low | CI.high | SE | Year | pred. | CI.low | CI.high | SE | Year | pred. | CI.low | CI.high | SE | Year | pred. | CI.low | CI.high | SE |
| Brownlees | 13 | 0.79 | 0.16 | 3.97 | 0.62 | 13 | 0.44 | 0.11 | 0.83 | 0.71 | 13 | 0.29 | 0.05 | 1.80 | 0.71 | 13 | 0.21 | 0.03 | 0.71 | 0.85 |
| Brownlees | 14 | 0.56 | 0.11 | 2.74 | 0.62 | 14 | 0.35 | 0.08 | 0.76 | 0.70 | 14 | 0.20 | 0.03 | 1.21 | 0.70 | 14 | 0.14 | 0.02 | 0.60 | 0.85 |
| Brownlees | 15 | 0.39 | 0.08 | 1.90 | 0.62 | 15 | 0.26 | 0.06 | 0.68 | 0.69 | 15 | 0.13 | 0.02 | 0.82 | 0.70 | 15 | 0.09 | 0.01 | 0.48 | 0.85 |
| Brownlees | 16 | 0.27 | 0.06 | 1.33 | 0.61 | 16 | 0.19 | 0.04 | 0.58 | 0.69 | 16 | 0.09 | 0.01 | 0.55 | 0.70 | 16 | 0.06 | 0.01 | 0.37 | 0.85 |
| Brownlees | 17 | 0.19 | 0.04 | 0.93 | 0.61 | 17 | 0.14 | 0.03 | 0.48 | 0.69 | 17 | 0.06 | 0.01 | 0.38 | 0.70 | 17 | 0.04 | 0.00 | 0.27 | 0.85 |
| Brownlees | 18 | 0.13 | 0.03 | 0.66 | 0.62 | 18 | 0.09 | 0.02 | 0.38 | 0.70 | 18 | 0.04 | 0.01 | 0.26 | 0.71 | 18 | 0.02 | 0.00 | 0.19 | 0.86 |
| Brownlees | 19 | 0.09 | 0.02 | 0.47 | 0.62 | 19 | 0.06 | 0.01 | 0.30 | 0.70 | 19 | 0.03 | 0.00 | 0.18 | 0.71 | 19 | 0.02 | 0.00 | 0.13 | 0.87 |
| Brownlees | 20 | 0.07 | 0.01 | 0.33 | 0.63 | 20 | 0.04 | 0.01 | 0.22 | 0.72 | 20 | 0.02 | 0.00 | 0.12 | 0.72 | 20 | 0.01 | 0.00 | 0.09 | 0.88 |
| Brownlees | 21 | 0.05 | 0.01 | 0.24 | 0.64 | 21 | 0.03 | 0.00 | 0.17 | 0.73 | 21 | 0.01 | 0.00 | 0.08 | 0.73 | 21 | 0.01 | 0.00 | 0.06 | 0.90 |
| Brownlees | 22 | 0.03 | 0.01 | 0.17 | 0.65 | 22 | 0.02 | 0.00 | 0.12 | 0.75 | 22 | 0.01 | 0.00 | 0.06 | 0.74 | 22 | 0.00 | 0.00 | 0.04 | 0.92 |
| Brownlees | 23 | 0.02 | 0.00 | 0.13 | 0.66 | 23 | 0.01 | 0.00 | 0.09 | 0.77 | 23 | 0.01 | 0.00 | 0.04 | 0.75 | 23 | 0.00 | 0.00 | 0.03 | 0.94 |
| Brownlees | 24 | 0.02 | 0.00 | 0.09 | 0.68 | 24 | 0.01 | 0.00 | 0.07 | 0.80 | 24 | 0.00 | 0.00 | 0.03 | 0.77 | 24 | 0.00 | 0.00 | 0.02 | 0.96 |
| Brownlees | 25 | 0.01 | 0.00 | 0.07 | 0.69 | 25 | 0.01 | 0.00 | 0.05 | 0.82 | 25 | 0.00 | 0.00 | 0.02 | 0.78 | 25 | 0.00 | 0.00 | 0.01 | 0.99 |
| Brownlees | 26 | 0.01 | 0.00 | 0.05 | 0.71 | 26 | 0.00 | 0.00 | 0.03 | 0.85 | 26 | 0.00 | 0.00 | 0.01 | 0.80 | 26 | 0.00 | 0.00 | 0.01 | 1.02 |
| Brownlees | 27 | 0.01 | 0.00 | 0.04 | 0.73 | 27 | 0.00 | 0.00 | 0.03 | 0.88 | 27 | 0.00 | 0.00 | 0.01 | 0.82 | 27 | 0.00 | 0.00 | 0.01 | 1.05 |
| Brownlees | 28 | 0.00 | 0.00 | 0.03 | 0.75 | 28 | 0.00 | 0.00 | 0.02 | 0.92 | 28 | 0.00 | 0.00 | 0.01 | 0.83 | 28 | 0.00 | 0.00 | 0.00 | 1.08 |
| Brownlees | 29 | 0.00 | 0.00 | 0.02 | 0.77 | 29 | 0.00 | 0.00 | 0.01 | 0.95 | 29 | 0.00 | 0.00 | 0.01 | 0.85 | 29 | 0.00 | 0.00 | 0.00 | 1.11 |
| Brownlees | 30 | 0.00 | 0.00 | 0.01 | 0.80 | 30 | 0.00 | 0.00 | 0.01 | 0.99 | 30 | 0.00 | 0.00 | 0.00 | 0.88 | 30 | 0.00 | 0.00 | 0.00 | 1.15 |
| Brownlees | 31 | 0.00 | 0.00 | 0.01 | 0.82 | 31 | 0.00 | 0.00 | 0.01 | 1.03 | 31 | 0.00 | 0.00 | 0.00 | 0.90 | 31 | 0.00 | 0.00 | 0.00 | 1.18 |
| Brownlees | 32 | 0.00 | 0.00 | 0.01 | 0.84 | 32 | 0.00 | 0.00 | 0.01 | 1.07 | 32 | 0.00 | 0.00 | 0.00 | 0.92 | 32 | 0.00 | 0.00 | 0.00 | 1.22 |
| Brownlees | 33 | 0.00 | 0.00 | 0.01 | 0.87 | 33 | 0.00 | 0.00 | 0.00 | 1.11 | 33 | 0.00 | 0.00 | 0.00 | 0.94 | 33 | 0.00 | 0.00 | 0.00 | 1.26 |
| Brownlees | 34 | 0.00 | 0.00 | 0.00 | 0.90 | 34 | 0.00 | 0.00 | 0.00 | 1.15 | 34 | 0.00 | 0.00 | 0.00 | 0.97 | 34 | 0.00 | 0.00 | 0.00 | 1.29 |
| Brownlees | 35 | 0.00 | 0.00 | 0.00 | 0.92 | 35 | 0.00 | 0.00 | 0.00 | 1.19 | 35 | 0.00 | 0.00 | 0.00 | 0.99 | 35 | 0.00 | 0.00 | 0.00 | 1.33 |
| Brownlees | 36 | 0.00 | 0.00 | 0.00 | 0.95 | 36 | 0.00 | 0.00 | 0.00 | 1.23 | 36 | 0.00 | 0.00 | 0.00 | 1.02 | 36 | 0.00 | 0.00 | 0.00 | 1.37 |

| | | | | | | | | | | | | | | | | | | | | |
|---|---|---|---|---|---|---|---|---|---|---|---|---|---|---|---|---|---|---|---|---|
| Brownlees | 37 | 0.00 | 0.00 | 0.00 | 0.98 | 37 | 0.00 | 0.00 | 0.00 | 1.27 | 37 | 0.00 | 0.00 | 0.00 | 1.04 | 37 | 0.00 | 0.00 | 0.00 | 1.42 |
| Brownlees | 38 | 0.00 | 0.00 | 0.00 | 1.01 | 38 | 0.00 | 0.00 | 0.00 | 1.32 | 38 | 0.00 | 0.00 | 0.00 | 1.07 | 38 | 0.00 | 0.00 | 0.00 | 1.46 |
| Brownlees | 39 | 0.00 | 0.00 | 0.00 | 1.04 | 39 | 0.00 | 0.00 | 0.00 | 1.36 | 39 | 0.00 | 0.00 | 0.00 | 1.10 | 39 | 0.00 | 0.00 | 0.00 | 1.50 |
| Brownlees | 40 | 0.00 | 0.00 | 0.00 | 1.07 | 40 | 0.00 | 0.00 | 0.00 | 1.41 | 40 | 0.00 | 0.00 | 0.00 | 1.13 | 40 | 0.00 | 0.00 | 0.00 | 1.54 |
| In_sounds | 13 | 0.46 | 0.14 | 1.51 | 0.46 | 13 | 0.41 | 0.17 | 0.70 | 0.48 | 13 | 0.27 | 0.06 | 1.19 | 0.57 | 13 | 0.26 | 0.06 | 0.66 | 0.65 |
| In_sounds | 14 | 0.32 | 0.10 | 1.05 | 0.46 | 14 | 0.31 | 0.12 | 0.60 | 0.47 | 14 | 0.18 | 0.04 | 0.80 | 0.57 | 14 | 0.18 | 0.04 | 0.54 | 0.65 |
| In_sounds | 15 | 0.23 | 0.07 | 0.73 | 0.46 | 15 | 0.23 | 0.08 | 0.50 | 0.46 | 15 | 0.12 | 0.03 | 0.54 | 0.57 | 15 | 0.12 | 0.03 | 0.42 | 0.64 |
| In_sounds | 16 | 0.16 | 0.05 | 0.52 | 0.46 | 16 | 0.17 | 0.06 | 0.40 | 0.46 | 16 | 0.08 | 0.02 | 0.37 | 0.57 | 16 | 0.08 | 0.02 | 0.31 | 0.64 |
| In_sounds | 17 | 0.11 | 0.03 | 0.37 | 0.46 | 17 | 0.12 | 0.04 | 0.31 | 0.47 | 17 | 0.06 | 0.01 | 0.25 | 0.57 | 17 | 0.05 | 0.01 | 0.22 | 0.65 |
| In_sounds | 18 | 0.08 | 0.02 | 0.26 | 0.47 | 18 | 0.08 | 0.03 | 0.24 | 0.48 | 18 | 0.04 | 0.01 | 0.17 | 0.58 | 18 | 0.03 | 0.01 | 0.15 | 0.66 |
| In_sounds | 19 | 0.05 | 0.02 | 0.19 | 0.48 | 19 | 0.06 | 0.02 | 0.18 | 0.50 | 19 | 0.03 | 0.01 | 0.12 | 0.59 | 19 | 0.02 | 0.00 | 0.11 | 0.67 |
| In_sounds | 20 | 0.04 | 0.01 | 0.14 | 0.49 | 20 | 0.04 | 0.01 | 0.13 | 0.52 | 20 | 0.02 | 0.00 | 0.08 | 0.60 | 20 | 0.01 | 0.00 | 0.07 | 0.69 |
| In_sounds | 21 | 0.03 | 0.01 | 0.10 | 0.51 | 21 | 0.03 | 0.01 | 0.10 | 0.54 | 21 | 0.01 | 0.00 | 0.06 | 0.61 | 21 | 0.01 | 0.00 | 0.05 | 0.71 |
| In_sounds | 22 | 0.02 | 0.00 | 0.07 | 0.52 | 22 | 0.02 | 0.00 | 0.07 | 0.57 | 22 | 0.01 | 0.00 | 0.04 | 0.62 | 22 | 0.00 | 0.00 | 0.03 | 0.73 |
| In_sounds | 23 | 0.01 | 0.00 | 0.05 | 0.54 | 23 | 0.01 | 0.00 | 0.05 | 0.61 | 23 | 0.01 | 0.00 | 0.03 | 0.64 | 23 | 0.00 | 0.00 | 0.02 | 0.76 |
| In_sounds | 24 | 0.01 | 0.00 | 0.04 | 0.56 | 24 | 0.01 | 0.00 | 0.04 | 0.64 | 24 | 0.00 | 0.00 | 0.02 | 0.65 | 24 | 0.00 | 0.00 | 0.01 | 0.79 |
| In_sounds | 25 | 0.01 | 0.00 | 0.03 | 0.58 | 25 | 0.01 | 0.00 | 0.03 | 0.68 | 25 | 0.00 | 0.00 | 0.01 | 0.67 | 25 | 0.00 | 0.00 | 0.01 | 0.82 |
| In_sounds | 26 | 0.00 | 0.00 | 0.02 | 0.61 | 26 | 0.00 | 0.00 | 0.02 | 0.72 | 26 | 0.00 | 0.00 | 0.01 | 0.69 | 26 | 0.00 | 0.00 | 0.01 | 0.85 |
| In_sounds | 27 | 0.00 | 0.00 | 0.02 | 0.63 | 27 | 0.00 | 0.00 | 0.02 | 0.76 | 27 | 0.00 | 0.00 | 0.01 | 0.71 | 27 | 0.00 | 0.00 | 0.00 | 0.89 |
| In_sounds | 28 | 0.00 | 0.00 | 0.01 | 0.66 | 28 | 0.00 | 0.00 | 0.01 | 0.80 | 28 | 0.00 | 0.00 | 0.01 | 0.74 | 28 | 0.00 | 0.00 | 0.00 | 0.93 |
| In_sounds | 29 | 0.00 | 0.00 | 0.01 | 0.69 | 29 | 0.00 | 0.00 | 0.01 | 0.84 | 29 | 0.00 | 0.00 | 0.00 | 0.76 | 29 | 0.00 | 0.00 | 0.00 | 0.96 |
| In_sounds | 30 | 0.00 | 0.00 | 0.01 | 0.71 | 30 | 0.00 | 0.00 | 0.01 | 0.89 | 30 | 0.00 | 0.00 | 0.00 | 0.78 | 30 | 0.00 | 0.00 | 0.00 | 1.00 |
| In_sounds | 31 | 0.00 | 0.00 | 0.01 | 0.74 | 31 | 0.00 | 0.00 | 0.00 | 0.93 | 31 | 0.00 | 0.00 | 0.00 | 0.81 | 31 | 0.00 | 0.00 | 0.00 | 1.04 |
| In_sounds | 32 | 0.00 | 0.00 | 0.00 | 0.77 | 32 | 0.00 | 0.00 | 0.00 | 0.98 | 32 | 0.00 | 0.00 | 0.00 | 0.83 | 32 | 0.00 | 0.00 | 0.00 | 1.08 |
| In_sounds | 33 | 0.00 | 0.00 | 0.00 | 0.80 | 33 | 0.00 | 0.00 | 0.00 | 1.02 | 33 | 0.00 | 0.00 | 0.00 | 0.86 | 33 | 0.00 | 0.00 | 0.00 | 1.13 |
| In_sounds | 34 | 0.00 | 0.00 | 0.00 | 0.83 | 34 | 0.00 | 0.00 | 0.00 | 1.07 | 34 | 0.00 | 0.00 | 0.00 | 0.89 | 34 | 0.00 | 0.00 | 0.00 | 1.17 |
| In_sounds | 35 | 0.00 | 0.00 | 0.00 | 0.86 | 35 | 0.00 | 0.00 | 0.00 | 1.11 | 35 | 0.00 | 0.00 | 0.00 | 0.91 | 35 | 0.00 | 0.00 | 0.00 | 1.21 |
| In_sounds | 36 | 0.00 | 0.00 | 0.00 | 0.89 | 36 | 0.00 | 0.00 | 0.00 | 1.16 | 36 | 0.00 | 0.00 | 0.00 | 0.94 | 36 | 0.00 | 0.00 | 0.00 | 1.26 |
| In_sounds | 37 | 0.00 | 0.00 | 0.00 | 0.93 | 37 | 0.00 | 0.00 | 0.00 | 1.21 | 37 | 0.00 | 0.00 | 0.00 | 0.97 | 37 | 0.00 | 0.00 | 0.00 | 1.30 |
| In_sounds | 38 | 0.00 | 0.00 | 0.00 | 0.96 | 38 | 0.00 | 0.00 | 0.00 | 1.26 | 38 | 0.00 | 0.00 | 0.00 | 1.00 | 38 | 0.00 | 0.00 | 0.00 | 1.35 |
| In_sounds | 39 | 0.00 | 0.00 | 0.00 | 0.99 | 39 | 0.00 | 0.00 | 0.00 | 1.31 | 39 | 0.00 | 0.00 | 0.00 | 1.03 | 39 | 0.00 | 0.00 | 0.00 | 1.39 |

| | | | | | | | | | | | | | | | | | | | | |
|---|---|---|---|---|---|---|---|---|---|---|---|---|---|---|---|---|---|---|---|---|
| In_sounds | 40 | 0.00 | 0.00 | 0.00 | 1.02 | 40 | 0.00 | 0.00 | 0.00 | 1.35 | 40 | 0.00 | 0.00 | 0.00 | 1.06 | 40 | 0.00 | 0.00 | 0.00 | 1.44 |
| Kaikumera | 13 | 0.77 | 0.22 | 2.71 | 0.49 | 13 | 0.48 | 0.19 | 0.78 | 0.54 | 13 | 0.52 | 0.09 | 2.94 | 0.67 | 13 | 0.36 | 0.07 | 0.82 | 0.80 |
| Kaikumera | 14 | 0.54 | 0.15 | 1.88 | 0.48 | 14 | 0.38 | 0.13 | 0.70 | 0.53 | 14 | 0.35 | 0.06 | 1.97 | 0.67 | 14 | 0.26 | 0.04 | 0.74 | 0.80 |
| Kaikumera | 15 | 0.38 | 0.11 | 1.32 | 0.48 | 15 | 0.29 | 0.09 | 0.61 | 0.52 | 15 | 0.24 | 0.04 | 1.33 | 0.67 | 15 | 0.18 | 0.03 | 0.63 | 0.79 |
| Kaikumera | 16 | 0.27 | 0.08 | 0.93 | 0.49 | 16 | 0.21 | 0.07 | 0.51 | 0.52 | 16 | 0.16 | 0.03 | 0.90 | 0.67 | 16 | 0.12 | 0.02 | 0.52 | 0.79 |
| Kaikumera | 17 | 0.19 | 0.05 | 0.66 | 0.49 | 17 | 0.15 | 0.04 | 0.41 | 0.53 | 17 | 0.11 | 0.02 | 0.62 | 0.68 | 17 | 0.08 | 0.01 | 0.40 | 0.80 |
| Kaikumera | 18 | 0.13 | 0.04 | 0.47 | 0.50 | 18 | 0.11 | 0.03 | 0.32 | 0.54 | 18 | 0.07 | 0.01 | 0.42 | 0.68 | 18 | 0.05 | 0.01 | 0.30 | 0.80 |
| Kaikumera | 19 | 0.09 | 0.02 | 0.34 | 0.51 | 19 | 0.07 | 0.02 | 0.25 | 0.55 | 19 | 0.05 | 0.01 | 0.29 | 0.69 | 19 | 0.03 | 0.00 | 0.21 | 0.81 |
| Kaikumera | 20 | 0.06 | 0.02 | 0.24 | 0.52 | 20 | 0.05 | 0.01 | 0.18 | 0.57 | 20 | 0.03 | 0.01 | 0.20 | 0.69 | 20 | 0.02 | 0.00 | 0.15 | 0.83 |
| Kaikumera | 21 | 0.04 | 0.01 | 0.18 | 0.53 | 21 | 0.03 | 0.01 | 0.14 | 0.59 | 21 | 0.02 | 0.00 | 0.14 | 0.70 | 21 | 0.01 | 0.00 | 0.10 | 0.84 |
| Kaikumera | 22 | 0.03 | 0.01 | 0.13 | 0.55 | 22 | 0.02 | 0.00 | 0.10 | 0.62 | 22 | 0.02 | 0.00 | 0.10 | 0.72 | 22 | 0.01 | 0.00 | 0.07 | 0.86 |
| Kaikumera | 23 | 0.02 | 0.01 | 0.10 | 0.57 | 23 | 0.02 | 0.00 | 0.08 | 0.65 | 23 | 0.01 | 0.00 | 0.07 | 0.73 | 23 | 0.00 | 0.00 | 0.05 | 0.88 |
| Kaikumera | 24 | 0.02 | 0.00 | 0.07 | 0.59 | 24 | 0.01 | 0.00 | 0.06 | 0.68 | 24 | 0.01 | 0.00 | 0.05 | 0.74 | 24 | 0.00 | 0.00 | 0.03 | 0.91 |
| Kaikumera | 25 | 0.01 | 0.00 | 0.05 | 0.61 | 25 | 0.01 | 0.00 | 0.04 | 0.72 | 25 | 0.00 | 0.00 | 0.03 | 0.76 | 25 | 0.00 | 0.00 | 0.02 | 0.93 |
| Kaikumera | 26 | 0.01 | 0.00 | 0.04 | 0.63 | 26 | 0.00 | 0.00 | 0.03 | 0.76 | 26 | 0.00 | 0.00 | 0.02 | 0.78 | 26 | 0.00 | 0.00 | 0.01 | 0.96 |
| Kaikumera | 27 | 0.01 | 0.00 | 0.03 | 0.66 | 27 | 0.00 | 0.00 | 0.02 | 0.79 | 27 | 0.00 | 0.00 | 0.02 | 0.80 | 27 | 0.00 | 0.00 | 0.01 | 0.99 |
| Kaikumera | 28 | 0.00 | 0.00 | 0.02 | 0.68 | 28 | 0.00 | 0.00 | 0.02 | 0.83 | 28 | 0.00 | 0.00 | 0.01 | 0.82 | 28 | 0.00 | 0.00 | 0.01 | 1.02 |
| Kaikumera | 29 | 0.00 | 0.00 | 0.02 | 0.71 | 29 | 0.00 | 0.00 | 0.01 | 0.87 | 29 | 0.00 | 0.00 | 0.01 | 0.84 | 29 | 0.00 | 0.00 | 0.00 | 1.06 |
| Kaikumera | 30 | 0.00 | 0.00 | 0.01 | 0.74 | 30 | 0.00 | 0.00 | 0.01 | 0.92 | 30 | 0.00 | 0.00 | 0.01 | 0.86 | 30 | 0.00 | 0.00 | 0.00 | 1.09 |
| Kaikumera | 31 | 0.00 | 0.00 | 0.01 | 0.76 | 31 | 0.00 | 0.00 | 0.01 | 0.96 | 31 | 0.00 | 0.00 | 0.00 | 0.88 | 31 | 0.00 | 0.00 | 0.00 | 1.13 |
| Kaikumera | 32 | 0.00 | 0.00 | 0.01 | 0.79 | 32 | 0.00 | 0.00 | 0.01 | 1.00 | 32 | 0.00 | 0.00 | 0.00 | 0.91 | 32 | 0.00 | 0.00 | 0.00 | 1.17 |
| Kaikumera | 33 | 0.00 | 0.00 | 0.01 | 0.82 | 33 | 0.00 | 0.00 | 0.00 | 1.05 | 33 | 0.00 | 0.00 | 0.00 | 0.93 | 33 | 0.00 | 0.00 | 0.00 | 1.20 |
| Kaikumera | 34 | 0.00 | 0.00 | 0.00 | 0.85 | 34 | 0.00 | 0.00 | 0.00 | 1.09 | 34 | 0.00 | 0.00 | 0.00 | 0.95 | 34 | 0.00 | 0.00 | 0.00 | 1.24 |
| Kaikumera | 35 | 0.00 | 0.00 | 0.00 | 0.88 | 35 | 0.00 | 0.00 | 0.00 | 1.14 | 35 | 0.00 | 0.00 | 0.00 | 0.98 | 35 | 0.00 | 0.00 | 0.00 | 1.28 |
| Kaikumera | 36 | 0.00 | 0.00 | 0.00 | 0.91 | 36 | 0.00 | 0.00 | 0.00 | 1.19 | 36 | 0.00 | 0.00 | 0.00 | 1.01 | 36 | 0.00 | 0.00 | 0.00 | 1.33 |
| Kaikumera | 37 | 0.00 | 0.00 | 0.00 | 0.94 | 37 | 0.00 | 0.00 | 0.00 | 1.23 | 37 | 0.00 | 0.00 | 0.00 | 1.03 | 37 | 0.00 | 0.00 | 0.00 | 1.37 |
| Kaikumera | 38 | 0.00 | 0.00 | 0.00 | 0.98 | 38 | 0.00 | 0.00 | 0.00 | 1.28 | 38 | 0.00 | 0.00 | 0.00 | 1.06 | 38 | 0.00 | 0.00 | 0.00 | 1.41 |
| Kaikumera | 39 | 0.00 | 0.00 | 0.00 | 1.01 | 39 | 0.00 | 0.00 | 0.00 | 1.33 | 39 | 0.00 | 0.00 | 0.00 | 1.09 | 39 | 0.00 | 0.00 | 0.00 | 1.45 |
| Kaikumera | 40 | 0.00 | 0.00 | 0.00 | 1.04 | 40 | 0.00 | 0.00 | 0.00 | 1.37 | 40 | 0.00 | 0.00 | 0.00 | 1.12 | 40 | 0.00 | 0.00 | 0.00 | 1.50 |
| Kaituna | 13 | 1.81 | 0.74 | 4.41 | 0.35 | 13 | 0.80 | 0.58 | 0.92 | 0.40 | 13 | 1.50 | 0.43 | 5.24 | 0.49 | 13 | 0.77 | 0.42 | 0.94 | 0.60 |
| Kaituna | 14 | 1.27 | 0.53 | 3.04 | 0.34 | 14 | 0.73 | 0.50 | 0.88 | 0.38 | 14 | 1.02 | 0.29 | 3.51 | 0.48 | 14 | 0.68 | 0.32 | 0.91 | 0.59 |

| | | | | | | | | | | | | | | | | | | | | |
|---|---|---|---|---|---|---|---|---|---|---|---|---|---|---|---|---|---|---|---|---|
| Kaituna | 15 | 0.89 | 0.37 | 2.12 | 0.34 | 15 | 0.64 | 0.41 | 0.82 | 0.36 | 15 | 0.69 | 0.20 | 2.37 | 0.48 | 15 | 0.57 | 0.23 | 0.85 | 0.58 |
| Kaituna | 16 | 0.62 | 0.26 | 1.49 | 0.34 | 16 | 0.54 | 0.32 | 0.74 | 0.35 | 16 | 0.47 | 0.13 | 1.61 | 0.48 | 16 | 0.45 | 0.16 | 0.78 | 0.57 |
| Kaituna | 17 | 0.44 | 0.18 | 1.06 | 0.34 | 17 | 0.44 | 0.24 | 0.65 | 0.35 | 17 | 0.32 | 0.09 | 1.10 | 0.49 | 17 | 0.34 | 0.11 | 0.69 | 0.57 |
| Kaituna | 18 | 0.31 | 0.12 | 0.76 | 0.35 | 18 | 0.34 | 0.17 | 0.56 | 0.35 | 18 | 0.21 | 0.06 | 0.76 | 0.49 | 18 | 0.24 | 0.07 | 0.58 | 0.57 |
| Kaituna | 19 | 0.21 | 0.08 | 0.55 | 0.36 | 19 | 0.26 | 0.12 | 0.47 | 0.36 | 19 | 0.14 | 0.04 | 0.52 | 0.50 | 19 | 0.17 | 0.04 | 0.47 | 0.58 |
| Kaituna | 20 | 0.15 | 0.06 | 0.40 | 0.38 | 20 | 0.19 | 0.08 | 0.38 | 0.38 | 20 | 0.10 | 0.03 | 0.37 | 0.51 | 20 | 0.11 | 0.03 | 0.36 | 0.59 |
| Kaituna | 21 | 0.11 | 0.04 | 0.30 | 0.40 | 21 | 0.13 | 0.05 | 0.30 | 0.40 | 21 | 0.07 | 0.02 | 0.26 | 0.53 | 21 | 0.07 | 0.02 | 0.27 | 0.61 |
| Kaituna | 22 | 0.07 | 0.03 | 0.22 | 0.42 | 22 | 0.09 | 0.03 | 0.23 | 0.43 | 22 | 0.04 | 0.01 | 0.18 | 0.54 | 22 | 0.05 | 0.01 | 0.19 | 0.63 |
| Kaituna | 23 | 0.05 | 0.02 | 0.16 | 0.44 | 23 | 0.06 | 0.02 | 0.18 | 0.46 | 23 | 0.03 | 0.01 | 0.13 | 0.56 | 23 | 0.03 | 0.01 | 0.14 | 0.65 |
| Kaituna | 24 | 0.04 | 0.01 | 0.12 | 0.47 | 24 | 0.04 | 0.01 | 0.14 | 0.50 | 24 | 0.02 | 0.00 | 0.09 | 0.58 | 24 | 0.02 | 0.00 | 0.10 | 0.68 |
| Kaituna | 25 | 0.03 | 0.01 | 0.09 | 0.49 | 25 | 0.03 | 0.01 | 0.11 | 0.54 | 25 | 0.01 | 0.00 | 0.06 | 0.60 | 25 | 0.01 | 0.00 | 0.07 | 0.71 |
| Kaituna | 26 | 0.02 | 0.00 | 0.07 | 0.52 | 26 | 0.02 | 0.00 | 0.08 | 0.58 | 26 | 0.01 | 0.00 | 0.05 | 0.62 | 26 | 0.01 | 0.00 | 0.05 | 0.74 |
| Kaituna | 27 | 0.01 | 0.00 | 0.05 | 0.55 | 27 | 0.01 | 0.00 | 0.06 | 0.62 | 27 | 0.01 | 0.00 | 0.03 | 0.64 | 27 | 0.00 | 0.00 | 0.03 | 0.78 |
| Kaituna | 28 | 0.01 | 0.00 | 0.04 | 0.58 | 28 | 0.01 | 0.00 | 0.05 | 0.66 | 28 | 0.00 | 0.00 | 0.02 | 0.67 | 28 | 0.00 | 0.00 | 0.02 | 0.81 |
| Kaituna | 29 | 0.01 | 0.00 | 0.03 | 0.61 | 29 | 0.01 | 0.00 | 0.03 | 0.71 | 29 | 0.00 | 0.00 | 0.02 | 0.69 | 29 | 0.00 | 0.00 | 0.02 | 0.85 |
| Kaituna | 30 | 0.00 | 0.00 | 0.02 | 0.64 | 30 | 0.00 | 0.00 | 0.03 | 0.75 | 30 | 0.00 | 0.00 | 0.01 | 0.72 | 30 | 0.00 | 0.00 | 0.01 | 0.89 |
| Kaituna | 31 | 0.00 | 0.00 | 0.02 | 0.67 | 31 | 0.00 | 0.00 | 0.02 | 0.80 | 31 | 0.00 | 0.00 | 0.01 | 0.75 | 31 | 0.00 | 0.00 | 0.01 | 0.93 |
| Kaituna | 32 | 0.00 | 0.00 | 0.01 | 0.70 | 32 | 0.00 | 0.00 | 0.01 | 0.85 | 32 | 0.00 | 0.00 | 0.01 | 0.77 | 32 | 0.00 | 0.00 | 0.00 | 0.97 |
| Kaituna | 33 | 0.00 | 0.00 | 0.01 | 0.74 | 33 | 0.00 | 0.00 | 0.01 | 0.89 | 33 | 0.00 | 0.00 | 0.00 | 0.80 | 33 | 0.00 | 0.00 | 0.00 | 1.01 |
| Kaituna | 34 | 0.00 | 0.00 | 0.01 | 0.77 | 34 | 0.00 | 0.00 | 0.01 | 0.94 | 34 | 0.00 | 0.00 | 0.00 | 0.83 | 34 | 0.00 | 0.00 | 0.00 | 1.06 |
| Kaituna | 35 | 0.00 | 0.00 | 0.01 | 0.80 | 35 | 0.00 | 0.00 | 0.01 | 0.99 | 35 | 0.00 | 0.00 | 0.00 | 0.86 | 35 | 0.00 | 0.00 | 0.00 | 1.10 |
| Kaituna | 36 | 0.00 | 0.00 | 0.00 | 0.84 | 36 | 0.00 | 0.00 | 0.00 | 1.04 | 36 | 0.00 | 0.00 | 0.00 | 0.89 | 36 | 0.00 | 0.00 | 0.00 | 1.14 |
| Kaituna | 37 | 0.00 | 0.00 | 0.00 | 0.87 | 37 | 0.00 | 0.00 | 0.00 | 1.09 | 37 | 0.00 | 0.00 | 0.00 | 0.92 | 37 | 0.00 | 0.00 | 0.00 | 1.19 |
| Kaituna | 38 | 0.00 | 0.00 | 0.00 | 0.90 | 38 | 0.00 | 0.00 | 0.00 | 1.14 | 38 | 0.00 | 0.00 | 0.00 | 0.95 | 38 | 0.00 | 0.00 | 0.00 | 1.24 |
| Kaituna | 39 | 0.00 | 0.00 | 0.00 | 0.94 | 39 | 0.00 | 0.00 | 0.00 | 1.19 | 39 | 0.00 | 0.00 | 0.00 | 0.98 | 39 | 0.00 | 0.00 | 0.00 | 1.28 |
| Kaituna | 40 | 0.00 | 0.00 | 0.00 | 0.97 | 40 | 0.00 | 0.00 | 0.00 | 1.24 | 40 | 0.00 | 0.00 | 0.00 | 1.01 | 40 | 0.00 | 0.00 | 0.00 | 1.33 |
| Kaiuma | 13 | 1.09 | 0.36 | 3.36 | 0.44 | 13 | 0.66 | 0.37 | 0.86 | 0.46 | 13 | 1.03 | 0.22 | 4.93 | 0.61 | 13 | 0.66 | 0.23 | 0.93 | 0.73 |
| Kaiuma | 14 | 0.77 | 0.25 | 2.33 | 0.43 | 14 | 0.56 | 0.29 | 0.80 | 0.45 | 14 | 0.70 | 0.15 | 3.31 | 0.60 | 14 | 0.55 | 0.16 | 0.88 | 0.72 |
| Kaiuma | 15 | 0.54 | 0.18 | 1.63 | 0.43 | 15 | 0.46 | 0.22 | 0.72 | 0.44 | 15 | 0.47 | 0.10 | 2.23 | 0.60 | 15 | 0.43 | 0.11 | 0.82 | 0.71 |
| Kaiuma | 16 | 0.38 | 0.12 | 1.15 | 0.43 | 16 | 0.36 | 0.16 | 0.63 | 0.43 | 16 | 0.32 | 0.07 | 1.52 | 0.60 | 16 | 0.32 | 0.07 | 0.74 | 0.71 |
| Kaiuma | 17 | 0.26 | 0.09 | 0.81 | 0.44 | 17 | 0.27 | 0.11 | 0.54 | 0.44 | 17 | 0.22 | 0.05 | 1.03 | 0.61 | 17 | 0.22 | 0.04 | 0.64 | 0.71 |

| | | | | | | | | | | | | | | | | | | | | |
|---|---|---|---|---|---|---|---|---|---|---|---|---|---|---|---|---|---|---|---|---|
| Kaiuma | 18 | 0.19 | 0.06 | 0.58 | 0.44 | 18 | 0.20 | 0.07 | 0.44 | 0.45 | 18 | 0.15 | 0.03 | 0.71 | 0.61 | 18 | 0.15 | 0.03 | 0.53 | 0.71 |
| Kaiuma | 19 | 0.13 | 0.04 | 0.42 | 0.46 | 19 | 0.14 | 0.05 | 0.35 | 0.46 | 19 | 0.10 | 0.02 | 0.49 | 0.62 | 19 | 0.10 | 0.02 | 0.42 | 0.72 |
| Kaiuma | 20 | 0.09 | 0.03 | 0.30 | 0.47 | 20 | 0.10 | 0.03 | 0.28 | 0.48 | 20 | 0.07 | 0.01 | 0.34 | 0.63 | 20 | 0.06 | 0.01 | 0.32 | 0.74 |
| Kaiuma | 21 | 0.06 | 0.02 | 0.22 | 0.48 | 21 | 0.07 | 0.02 | 0.21 | 0.51 | 21 | 0.05 | 0.01 | 0.24 | 0.64 | 21 | 0.04 | 0.01 | 0.23 | 0.75 |
| Kaiuma | 22 | 0.04 | 0.01 | 0.16 | 0.50 | 22 | 0.05 | 0.01 | 0.16 | 0.53 | 22 | 0.03 | 0.01 | 0.17 | 0.65 | 22 | 0.03 | 0.00 | 0.16 | 0.77 |
| Kaiuma | 23 | 0.03 | 0.01 | 0.12 | 0.52 | 23 | 0.03 | 0.01 | 0.12 | 0.57 | 23 | 0.02 | 0.00 | 0.12 | 0.67 | 23 | 0.02 | 0.00 | 0.11 | 0.79 |
| Kaiuma | 24 | 0.02 | 0.01 | 0.09 | 0.54 | 24 | 0.02 | 0.00 | 0.09 | 0.60 | 24 | 0.01 | 0.00 | 0.08 | 0.68 | 24 | 0.01 | 0.00 | 0.08 | 0.82 |
| Kaiuma | 25 | 0.02 | 0.00 | 0.07 | 0.57 | 25 | 0.01 | 0.00 | 0.07 | 0.64 | 25 | 0.01 | 0.00 | 0.06 | 0.70 | 25 | 0.01 | 0.00 | 0.05 | 0.84 |
| Kaiuma | 26 | 0.01 | 0.00 | 0.05 | 0.59 | 26 | 0.01 | 0.00 | 0.05 | 0.68 | 26 | 0.01 | 0.00 | 0.04 | 0.72 | 26 | 0.00 | 0.00 | 0.04 | 0.87 |
| Kaiuma | 27 | 0.01 | 0.00 | 0.04 | 0.62 | 27 | 0.01 | 0.00 | 0.04 | 0.72 | 27 | 0.00 | 0.00 | 0.03 | 0.74 | 27 | 0.00 | 0.00 | 0.02 | 0.90 |
| Kaiuma | 28 | 0.01 | 0.00 | 0.03 | 0.64 | 28 | 0.00 | 0.00 | 0.03 | 0.76 | 28 | 0.00 | 0.00 | 0.02 | 0.76 | 28 | 0.00 | 0.00 | 0.02 | 0.94 |
| Kaiuma | 29 | 0.00 | 0.00 | 0.02 | 0.67 | 29 | 0.00 | 0.00 | 0.02 | 0.80 | 29 | 0.00 | 0.00 | 0.02 | 0.78 | 29 | 0.00 | 0.00 | 0.01 | 0.97 |
| Kaiuma | 30 | 0.00 | 0.00 | 0.02 | 0.70 | 30 | 0.00 | 0.00 | 0.02 | 0.85 | 30 | 0.00 | 0.00 | 0.01 | 0.81 | 30 | 0.00 | 0.00 | 0.01 | 1.01 |
| Kaiuma | 31 | 0.00 | 0.00 | 0.01 | 0.73 | 31 | 0.00 | 0.00 | 0.01 | 0.89 | 31 | 0.00 | 0.00 | 0.01 | 0.83 | 31 | 0.00 | 0.00 | 0.01 | 1.04 |
| Kaiuma | 32 | 0.00 | 0.00 | 0.01 | 0.76 | 32 | 0.00 | 0.00 | 0.01 | 0.94 | 32 | 0.00 | 0.00 | 0.01 | 0.86 | 32 | 0.00 | 0.00 | 0.00 | 1.08 |
| Kaiuma | 33 | 0.00 | 0.00 | 0.01 | 0.79 | 33 | 0.00 | 0.00 | 0.01 | 0.98 | 33 | 0.00 | 0.00 | 0.00 | 0.88 | 33 | 0.00 | 0.00 | 0.00 | 1.12 |
| Kaiuma | 34 | 0.00 | 0.00 | 0.01 | 0.82 | 34 | 0.00 | 0.00 | 0.01 | 1.03 | 34 | 0.00 | 0.00 | 0.00 | 0.91 | 34 | 0.00 | 0.00 | 0.00 | 1.16 |
| Kaiuma | 35 | 0.00 | 0.00 | 0.00 | 0.85 | 35 | 0.00 | 0.00 | 0.00 | 1.08 | 35 | 0.00 | 0.00 | 0.00 | 0.93 | 35 | 0.00 | 0.00 | 0.00 | 1.20 |
| Kaiuma | 36 | 0.00 | 0.00 | 0.00 | 0.88 | 36 | 0.00 | 0.00 | 0.00 | 1.12 | 36 | 0.00 | 0.00 | 0.00 | 0.96 | 36 | 0.00 | 0.00 | 0.00 | 1.25 |
| Kaiuma | 37 | 0.00 | 0.00 | 0.00 | 0.92 | 37 | 0.00 | 0.00 | 0.00 | 1.17 | 37 | 0.00 | 0.00 | 0.00 | 0.99 | 37 | 0.00 | 0.00 | 0.00 | 1.29 |
| Kaiuma | 38 | 0.00 | 0.00 | 0.00 | 0.95 | 38 | 0.00 | 0.00 | 0.00 | 1.22 | 38 | 0.00 | 0.00 | 0.00 | 1.02 | 38 | 0.00 | 0.00 | 0.00 | 1.33 |
| Kaiuma | 39 | 0.00 | 0.00 | 0.00 | 0.98 | 39 | 0.00 | 0.00 | 0.00 | 1.27 | 39 | 0.00 | 0.00 | 0.00 | 1.05 | 39 | 0.00 | 0.00 | 0.00 | 1.38 |
| Kaiuma | 40 | 0.00 | 0.00 | 0.00 | 1.02 | 40 | 0.00 | 0.00 | 0.00 | 1.32 | 40 | 0.00 | 0.00 | 0.00 | 1.08 | 40 | 0.00 | 0.00 | 0.00 | 1.42 |
| Kenepuru | 13 | 0.27 | 0.07 | 1.12 | 0.55 | 13 | 0.32 | 0.11 | 0.65 | 0.54 | 13 | 0.14 | 0.03 | 0.75 | 0.64 | 13 | 0.17 | 0.03 | 0.54 | 0.68 |
| Kenepuru | 14 | 0.19 | 0.05 | 0.78 | 0.54 | 14 | 0.24 | 0.07 | 0.55 | 0.53 | 14 | 0.10 | 0.02 | 0.51 | 0.64 | 14 | 0.11 | 0.02 | 0.42 | 0.68 |
| Kenepuru | 15 | 0.13 | 0.03 | 0.55 | 0.54 | 15 | 0.17 | 0.05 | 0.45 | 0.53 | 15 | 0.07 | 0.01 | 0.34 | 0.64 | 15 | 0.07 | 0.01 | 0.31 | 0.67 |
| Kenepuru | 16 | 0.09 | 0.02 | 0.39 | 0.55 | 16 | 0.12 | 0.03 | 0.35 | 0.53 | 16 | 0.04 | 0.01 | 0.23 | 0.64 | 16 | 0.05 | 0.01 | 0.22 | 0.68 |
| Kenepuru | 17 | 0.07 | 0.02 | 0.27 | 0.55 | 17 | 0.08 | 0.02 | 0.27 | 0.54 | 17 | 0.03 | 0.01 | 0.16 | 0.64 | 17 | 0.03 | 0.01 | 0.15 | 0.68 |
| Kenepuru | 18 | 0.05 | 0.01 | 0.19 | 0.56 | 18 | 0.06 | 0.01 | 0.20 | 0.55 | 18 | 0.02 | 0.00 | 0.11 | 0.65 | 18 | 0.02 | 0.00 | 0.10 | 0.69 |
| Kenepuru | 19 | 0.03 | 0.01 | 0.14 | 0.57 | 19 | 0.04 | 0.01 | 0.15 | 0.57 | 19 | 0.01 | 0.00 | 0.07 | 0.65 | 19 | 0.01 | 0.00 | 0.07 | 0.71 |
| Kenepuru | 20 | 0.02 | 0.01 | 0.10 | 0.58 | 20 | 0.03 | 0.01 | 0.11 | 0.59 | 20 | 0.01 | 0.00 | 0.05 | 0.66 | 20 | 0.01 | 0.00 | 0.05 | 0.72 |

| | | | | | | | | | | | | | | | | | | | | |
|---|---|---|---|---|---|---|---|---|---|---|---|---|---|---|---|---|---|---|---|---|
| Kenepuru | 21 | 0.02 | 0.00 | 0.07 | 0.59 | 21 | 0.02 | 0.00 | 0.08 | 0.61 | 21 | 0.01 | 0.00 | 0.04 | 0.67 | 21 | 0.00 | 0.00 | 0.03 | 0.75 |
| Kenepuru | 22 | 0.01 | 0.00 | 0.05 | 0.61 | 22 | 0.01 | 0.00 | 0.06 | 0.64 | 22 | 0.00 | 0.00 | 0.03 | 0.69 | 22 | 0.00 | 0.00 | 0.02 | 0.77 |
| Kenepuru | 23 | 0.01 | 0.00 | 0.04 | 0.62 | 23 | 0.01 | 0.00 | 0.04 | 0.67 | 23 | 0.00 | 0.00 | 0.02 | 0.70 | 23 | 0.00 | 0.00 | 0.01 | 0.79 |
| Kenepuru | 24 | 0.01 | 0.00 | 0.03 | 0.64 | 24 | 0.01 | 0.00 | 0.03 | 0.70 | 24 | 0.00 | 0.00 | 0.01 | 0.72 | 24 | 0.00 | 0.00 | 0.01 | 0.82 |
| Kenepuru | 25 | 0.00 | 0.00 | 0.02 | 0.66 | 25 | 0.00 | 0.00 | 0.02 | 0.74 | 25 | 0.00 | 0.00 | 0.01 | 0.73 | 25 | 0.00 | 0.00 | 0.01 | 0.85 |
| Kenepuru | 26 | 0.00 | 0.00 | 0.02 | 0.68 | 26 | 0.00 | 0.00 | 0.02 | 0.78 | 26 | 0.00 | 0.00 | 0.01 | 0.75 | 26 | 0.00 | 0.00 | 0.00 | 0.89 |
| Kenepuru | 27 | 0.00 | 0.00 | 0.01 | 0.71 | 27 | 0.00 | 0.00 | 0.01 | 0.81 | 27 | 0.00 | 0.00 | 0.00 | 0.77 | 27 | 0.00 | 0.00 | 0.00 | 0.92 |
| Kenepuru | 28 | 0.00 | 0.00 | 0.01 | 0.73 | 28 | 0.00 | 0.00 | 0.01 | 0.85 | 28 | 0.00 | 0.00 | 0.00 | 0.79 | 28 | 0.00 | 0.00 | 0.00 | 0.96 |
| Kenepuru | 29 | 0.00 | 0.00 | 0.01 | 0.76 | 29 | 0.00 | 0.00 | 0.01 | 0.90 | 29 | 0.00 | 0.00 | 0.00 | 0.81 | 29 | 0.00 | 0.00 | 0.00 | 1.00 |
| Kenepuru | 30 | 0.00 | 0.00 | 0.00 | 0.78 | 30 | 0.00 | 0.00 | 0.01 | 0.94 | 30 | 0.00 | 0.00 | 0.00 | 0.83 | 30 | 0.00 | 0.00 | 0.00 | 1.03 |
| Kenepuru | 31 | 0.00 | 0.00 | 0.00 | 0.81 | 31 | 0.00 | 0.00 | 0.00 | 0.98 | 31 | 0.00 | 0.00 | 0.00 | 0.86 | 31 | 0.00 | 0.00 | 0.00 | 1.07 |
| Kenepuru | 32 | 0.00 | 0.00 | 0.00 | 0.84 | 32 | 0.00 | 0.00 | 0.00 | 1.03 | 32 | 0.00 | 0.00 | 0.00 | 0.88 | 32 | 0.00 | 0.00 | 0.00 | 1.11 |
| Kenepuru | 33 | 0.00 | 0.00 | 0.00 | 0.86 | 33 | 0.00 | 0.00 | 0.00 | 1.07 | 33 | 0.00 | 0.00 | 0.00 | 0.91 | 33 | 0.00 | 0.00 | 0.00 | 1.16 |
| Kenepuru | 34 | 0.00 | 0.00 | 0.00 | 0.89 | 34 | 0.00 | 0.00 | 0.00 | 1.12 | 34 | 0.00 | 0.00 | 0.00 | 0.93 | 34 | 0.00 | 0.00 | 0.00 | 1.20 |
| Kenepuru | 35 | 0.00 | 0.00 | 0.00 | 0.92 | 35 | 0.00 | 0.00 | 0.00 | 1.16 | 35 | 0.00 | 0.00 | 0.00 | 0.96 | 35 | 0.00 | 0.00 | 0.00 | 1.24 |
| Kenepuru | 36 | 0.00 | 0.00 | 0.00 | 0.95 | 36 | 0.00 | 0.00 | 0.00 | 1.21 | 36 | 0.00 | 0.00 | 0.00 | 0.99 | 36 | 0.00 | 0.00 | 0.00 | 1.29 |
| Kenepuru | 37 | 0.00 | 0.00 | 0.00 | 0.98 | 37 | 0.00 | 0.00 | 0.00 | 1.25 | 37 | 0.00 | 0.00 | 0.00 | 1.01 | 37 | 0.00 | 0.00 | 0.00 | 1.33 |
| Kenepuru | 38 | 0.00 | 0.00 | 0.00 | 1.01 | 38 | 0.00 | 0.00 | 0.00 | 1.30 | 38 | 0.00 | 0.00 | 0.00 | 1.04 | 38 | 0.00 | 0.00 | 0.00 | 1.37 |
| Kenepuru | 39 | 0.00 | 0.00 | 0.00 | 1.04 | 39 | 0.00 | 0.00 | 0.00 | 1.35 | 39 | 0.00 | 0.00 | 0.00 | 1.07 | 39 | 0.00 | 0.00 | 0.00 | 1.42 |
| Kenepuru | 40 | 0.00 | 0.00 | 0.00 | 1.07 | 40 | 0.00 | 0.00 | 0.00 | 1.40 | 40 | 0.00 | 0.00 | 0.00 | 1.10 | 40 | 0.00 | 0.00 | 0.00 | 1.47 |
| Mahakipawa | 13 | 0.80 | 0.35 | 1.83 | 0.32 | 13 | 0.57 | 0.35 | 0.77 | 0.35 | 13 | 0.66 | 0.22 | 2.00 | 0.43 | 13 | 0.52 | 0.23 | 0.80 | 0.51 |
| Mahakipawa | 14 | 0.56 | 0.25 | 1.26 | 0.32 | 14 | 0.47 | 0.27 | 0.68 | 0.33 | 14 | 0.45 | 0.15 | 1.34 | 0.43 | 14 | 0.40 | 0.16 | 0.71 | 0.50 |
| Mahakipawa | 15 | 0.39 | 0.18 | 0.88 | 0.31 | 15 | 0.37 | 0.20 | 0.58 | 0.32 | 15 | 0.30 | 0.10 | 0.90 | 0.42 | 15 | 0.30 | 0.11 | 0.60 | 0.49 |
| Mahakipawa | 16 | 0.28 | 0.12 | 0.62 | 0.32 | 16 | 0.28 | 0.15 | 0.47 | 0.32 | 16 | 0.20 | 0.07 | 0.61 | 0.43 | 16 | 0.21 | 0.07 | 0.48 | 0.49 |
| Mahakipawa | 17 | 0.19 | 0.08 | 0.45 | 0.32 | 17 | 0.21 | 0.10 | 0.38 | 0.33 | 17 | 0.14 | 0.05 | 0.42 | 0.43 | 17 | 0.14 | 0.04 | 0.37 | 0.49 |
| Mahakipawa | 18 | 0.14 | 0.06 | 0.32 | 0.34 | 18 | 0.15 | 0.07 | 0.30 | 0.34 | 18 | 0.09 | 0.03 | 0.29 | 0.44 | 18 | 0.09 | 0.03 | 0.27 | 0.50 |
| Mahakipawa | 19 | 0.09 | 0.04 | 0.23 | 0.35 | 19 | 0.10 | 0.04 | 0.23 | 0.37 | 19 | 0.06 | 0.02 | 0.20 | 0.45 | 19 | 0.06 | 0.02 | 0.19 | 0.52 |
| Mahakipawa | 20 | 0.07 | 0.03 | 0.17 | 0.37 | 20 | 0.07 | 0.03 | 0.17 | 0.39 | 20 | 0.04 | 0.01 | 0.14 | 0.46 | 20 | 0.04 | 0.01 | 0.14 | 0.54 |
| Mahakipawa | 21 | 0.05 | 0.02 | 0.13 | 0.39 | 21 | 0.05 | 0.02 | 0.13 | 0.43 | 21 | 0.03 | 0.01 | 0.10 | 0.48 | 21 | 0.02 | 0.01 | 0.09 | 0.56 |
| Mahakipawa | 22 | 0.03 | 0.01 | 0.09 | 0.41 | 22 | 0.03 | 0.01 | 0.10 | 0.46 | 22 | 0.02 | 0.01 | 0.07 | 0.49 | 22 | 0.01 | 0.00 | 0.06 | 0.59 |
| Mahakipawa | 23 | 0.02 | 0.01 | 0.07 | 0.43 | 23 | 0.02 | 0.01 | 0.08 | 0.50 | 23 | 0.01 | 0.00 | 0.05 | 0.51 | 23 | 0.01 | 0.00 | 0.04 | 0.62 |

| | | | | | | | | | | | | | | | | | | | | |
|---|---|---|---|---|---|---|---|---|---|---|---|---|---|---|---|---|---|---|---|---|
| Mahakipawa | 24 | 0.02 | 0.00 | 0.05 | 0.46 | 24 | 0.01 | 0.00 | 0.06 | 0.54 | 24 | 0.01 | 0.00 | 0.04 | 0.53 | 24 | 0.01 | 0.00 | 0.03 | 0.65 |
| Mahakipawa | 25 | 0.01 | 0.00 | 0.04 | 0.49 | 25 | 0.01 | 0.00 | 0.04 | 0.58 | 25 | 0.01 | 0.00 | 0.03 | 0.56 | 25 | 0.00 | 0.00 | 0.02 | 0.69 |
| Mahakipawa | 26 | 0.01 | 0.00 | 0.03 | 0.52 | 26 | 0.01 | 0.00 | 0.03 | 0.63 | 26 | 0.00 | 0.00 | 0.02 | 0.58 | 26 | 0.00 | 0.00 | 0.01 | 0.72 |
| Mahakipawa | 27 | 0.01 | 0.00 | 0.02 | 0.55 | 27 | 0.00 | 0.00 | 0.02 | 0.67 | 27 | 0.00 | 0.00 | 0.01 | 0.60 | 27 | 0.00 | 0.00 | 0.01 | 0.76 |
| Mahakipawa | 28 | 0.00 | 0.00 | 0.02 | 0.58 | 28 | 0.00 | 0.00 | 0.02 | 0.72 | 28 | 0.00 | 0.00 | 0.01 | 0.63 | 28 | 0.00 | 0.00 | 0.01 | 0.80 |
| Mahakipawa | 29 | 0.00 | 0.00 | 0.01 | 0.61 | 29 | 0.00 | 0.00 | 0.01 | 0.76 | 29 | 0.00 | 0.00 | 0.01 | 0.66 | 29 | 0.00 | 0.00 | 0.00 | 0.85 |
| Mahakipawa | 30 | 0.00 | 0.00 | 0.01 | 0.64 | 30 | 0.00 | 0.00 | 0.01 | 0.81 | 30 | 0.00 | 0.00 | 0.01 | 0.68 | 30 | 0.00 | 0.00 | 0.00 | 0.89 |
| Mahakipawa | 31 | 0.00 | 0.00 | 0.01 | 0.67 | 31 | 0.00 | 0.00 | 0.01 | 0.86 | 31 | 0.00 | 0.00 | 0.00 | 0.71 | 31 | 0.00 | 0.00 | 0.00 | 0.93 |
| Mahakipawa | 32 | 0.00 | 0.00 | 0.01 | 0.70 | 32 | 0.00 | 0.00 | 0.01 | 0.91 | 32 | 0.00 | 0.00 | 0.00 | 0.74 | 32 | 0.00 | 0.00 | 0.00 | 0.98 |
| Mahakipawa | 33 | 0.00 | 0.00 | 0.00 | 0.74 | 33 | 0.00 | 0.00 | 0.00 | 0.96 | 33 | 0.00 | 0.00 | 0.00 | 0.77 | 33 | 0.00 | 0.00 | 0.00 | 1.02 |
| Mahakipawa | 34 | 0.00 | 0.00 | 0.00 | 0.77 | 34 | 0.00 | 0.00 | 0.00 | 1.01 | 34 | 0.00 | 0.00 | 0.00 | 0.80 | 34 | 0.00 | 0.00 | 0.00 | 1.07 |
| Mahakipawa | 35 | 0.00 | 0.00 | 0.00 | 0.80 | 35 | 0.00 | 0.00 | 0.00 | 1.05 | 35 | 0.00 | 0.00 | 0.00 | 0.83 | 35 | 0.00 | 0.00 | 0.00 | 1.11 |
| Mahakipawa | 36 | 0.00 | 0.00 | 0.00 | 0.84 | 36 | 0.00 | 0.00 | 0.00 | 1.10 | 36 | 0.00 | 0.00 | 0.00 | 0.86 | 36 | 0.00 | 0.00 | 0.00 | 1.16 |
| Mahakipawa | 37 | 0.00 | 0.00 | 0.00 | 0.87 | 37 | 0.00 | 0.00 | 0.00 | 1.15 | 37 | 0.00 | 0.00 | 0.00 | 0.89 | 37 | 0.00 | 0.00 | 0.00 | 1.21 |
| Mahakipawa | 38 | 0.00 | 0.00 | 0.00 | 0.91 | 38 | 0.00 | 0.00 | 0.00 | 1.20 | 38 | 0.00 | 0.00 | 0.00 | 0.92 | 38 | 0.00 | 0.00 | 0.00 | 1.26 |
| Mahakipawa | 39 | 0.00 | 0.00 | 0.00 | 0.94 | 39 | 0.00 | 0.00 | 0.00 | 1.25 | 39 | 0.00 | 0.00 | 0.00 | 0.96 | 39 | 0.00 | 0.00 | 0.00 | 1.30 |
| Mahakipawa | 40 | 0.00 | 0.00 | 0.00 | 0.98 | 40 | 0.00 | 0.00 | 0.00 | 1.30 | 40 | 0.00 | 0.00 | 0.00 | 0.99 | 40 | 0.00 | 0.00 | 0.00 | 1.35 |
| Mahau | 13 | 1.09 | 0.33 | 3.67 | 0.47 | 13 | 0.65 | 0.34 | 0.87 | 0.50 | 13 | 0.75 | 0.15 | 3.89 | 0.64 | 13 | 0.55 | 0.15 | 0.90 | 0.75 |
| Mahau | 14 | 0.77 | 0.23 | 2.54 | 0.47 | 14 | 0.55 | 0.26 | 0.81 | 0.49 | 14 | 0.51 | 0.10 | 2.61 | 0.63 | 14 | 0.44 | 0.10 | 0.84 | 0.75 |
| Mahau | 15 | 0.54 | 0.16 | 1.77 | 0.46 | 15 | 0.45 | 0.20 | 0.74 | 0.48 | 15 | 0.35 | 0.07 | 1.76 | 0.63 | 15 | 0.32 | 0.07 | 0.76 | 0.74 |
| Mahau | 16 | 0.38 | 0.11 | 1.25 | 0.46 | 16 | 0.36 | 0.14 | 0.65 | 0.47 | 16 | 0.23 | 0.05 | 1.20 | 0.63 | 16 | 0.23 | 0.04 | 0.67 | 0.74 |
| Mahau | 17 | 0.26 | 0.08 | 0.88 | 0.47 | 17 | 0.27 | 0.10 | 0.55 | 0.48 | 17 | 0.16 | 0.03 | 0.82 | 0.64 | 17 | 0.16 | 0.03 | 0.56 | 0.74 |
| Mahau | 18 | 0.19 | 0.05 | 0.63 | 0.47 | 18 | 0.20 | 0.07 | 0.46 | 0.48 | 18 | 0.11 | 0.02 | 0.56 | 0.64 | 18 | 0.10 | 0.02 | 0.44 | 0.75 |
| Mahau | 19 | 0.13 | 0.04 | 0.45 | 0.48 | 19 | 0.14 | 0.04 | 0.37 | 0.50 | 19 | 0.07 | 0.01 | 0.39 | 0.65 | 19 | 0.07 | 0.01 | 0.34 | 0.76 |
| Mahau | 20 | 0.09 | 0.03 | 0.33 | 0.50 | 20 | 0.10 | 0.03 | 0.29 | 0.51 | 20 | 0.05 | 0.01 | 0.27 | 0.66 | 20 | 0.04 | 0.01 | 0.25 | 0.77 |
| Mahau | 21 | 0.06 | 0.02 | 0.24 | 0.51 | 21 | 0.07 | 0.02 | 0.22 | 0.54 | 21 | 0.03 | 0.01 | 0.19 | 0.67 | 21 | 0.03 | 0.00 | 0.17 | 0.79 |
| Mahau | 22 | 0.04 | 0.01 | 0.17 | 0.53 | 22 | 0.05 | 0.01 | 0.17 | 0.56 | 22 | 0.02 | 0.00 | 0.13 | 0.68 | 22 | 0.02 | 0.00 | 0.12 | 0.80 |
| Mahau | 23 | 0.03 | 0.01 | 0.13 | 0.55 | 23 | 0.03 | 0.01 | 0.13 | 0.59 | 23 | 0.02 | 0.00 | 0.09 | 0.70 | 23 | 0.01 | 0.00 | 0.08 | 0.83 |
| Mahau | 24 | 0.02 | 0.01 | 0.09 | 0.57 | 24 | 0.02 | 0.00 | 0.09 | 0.62 | 24 | 0.01 | 0.00 | 0.06 | 0.71 | 24 | 0.01 | 0.00 | 0.06 | 0.85 |
| Mahau | 25 | 0.02 | 0.00 | 0.07 | 0.59 | 25 | 0.01 | 0.00 | 0.07 | 0.66 | 25 | 0.01 | 0.00 | 0.05 | 0.73 | 25 | 0.00 | 0.00 | 0.04 | 0.88 |
| Mahau | 26 | 0.01 | 0.00 | 0.05 | 0.61 | 26 | 0.01 | 0.00 | 0.05 | 0.70 | 26 | 0.00 | 0.00 | 0.03 | 0.75 | 26 | 0.00 | 0.00 | 0.03 | 0.91 |

| | | | | | | | | | | | | | | | | | | | | |
|---|---|---|---|---|---|---|---|---|---|---|---|---|---|---|---|---|---|---|---|---|
| Mahau | 27 | 0.01 | 0.00 | 0.04 | 0.64 | 27 | 0.01 | 0.00 | 0.04 | 0.73 | 27 | 0.00 | 0.00 | 0.02 | 0.77 | 27 | 0.00 | 0.00 | 0.02 | 0.94 |
| Mahau | 28 | 0.01 | 0.00 | 0.03 | 0.66 | 28 | 0.00 | 0.00 | 0.03 | 0.77 | 28 | 0.00 | 0.00 | 0.02 | 0.79 | 28 | 0.00 | 0.00 | 0.01 | 0.97 |
| Mahau | 29 | 0.00 | 0.00 | 0.02 | 0.69 | 29 | 0.00 | 0.00 | 0.02 | 0.82 | 29 | 0.00 | 0.00 | 0.01 | 0.81 | 29 | 0.00 | 0.00 | 0.01 | 1.00 |
| Mahau | 30 | 0.00 | 0.00 | 0.02 | 0.71 | 30 | 0.00 | 0.00 | 0.02 | 0.86 | 30 | 0.00 | 0.00 | 0.01 | 0.83 | 30 | 0.00 | 0.00 | 0.01 | 1.04 |
| Mahau | 31 | 0.00 | 0.00 | 0.01 | 0.74 | 31 | 0.00 | 0.00 | 0.01 | 0.90 | 31 | 0.00 | 0.00 | 0.01 | 0.85 | 31 | 0.00 | 0.00 | 0.00 | 1.08 |
| Mahau | 32 | 0.00 | 0.00 | 0.01 | 0.77 | 32 | 0.00 | 0.00 | 0.01 | 0.95 | 32 | 0.00 | 0.00 | 0.00 | 0.88 | 32 | 0.00 | 0.00 | 0.00 | 1.12 |
| Mahau | 33 | 0.00 | 0.00 | 0.01 | 0.80 | 33 | 0.00 | 0.00 | 0.01 | 0.99 | 33 | 0.00 | 0.00 | 0.00 | 0.90 | 33 | 0.00 | 0.00 | 0.00 | 1.16 |
| Mahau | 34 | 0.00 | 0.00 | 0.01 | 0.83 | 34 | 0.00 | 0.00 | 0.01 | 1.04 | 34 | 0.00 | 0.00 | 0.00 | 0.93 | 34 | 0.00 | 0.00 | 0.00 | 1.20 |
| Mahau | 35 | 0.00 | 0.00 | 0.00 | 0.86 | 35 | 0.00 | 0.00 | 0.00 | 1.08 | 35 | 0.00 | 0.00 | 0.00 | 0.96 | 35 | 0.00 | 0.00 | 0.00 | 1.24 |
| Mahau | 36 | 0.00 | 0.00 | 0.00 | 0.89 | 36 | 0.00 | 0.00 | 0.00 | 1.13 | 36 | 0.00 | 0.00 | 0.00 | 0.98 | 36 | 0.00 | 0.00 | 0.00 | 1.28 |
| Mahau | 37 | 0.00 | 0.00 | 0.00 | 0.93 | 37 | 0.00 | 0.00 | 0.00 | 1.18 | 37 | 0.00 | 0.00 | 0.00 | 1.01 | 37 | 0.00 | 0.00 | 0.00 | 1.32 |
| Mahau | 38 | 0.00 | 0.00 | 0.00 | 0.96 | 38 | 0.00 | 0.00 | 0.00 | 1.23 | 38 | 0.00 | 0.00 | 0.00 | 1.04 | 38 | 0.00 | 0.00 | 0.00 | 1.36 |
| Mahau | 39 | 0.00 | 0.00 | 0.00 | 0.99 | 39 | 0.00 | 0.00 | 0.00 | 1.27 | 39 | 0.00 | 0.00 | 0.00 | 1.07 | 39 | 0.00 | 0.00 | 0.00 | 1.41 |
| Mahau | 40 | 0.00 | 0.00 | 0.00 | 1.02 | 40 | 0.00 | 0.00 | 0.00 | 1.32 | 40 | 0.00 | 0.00 | 0.00 | 1.10 | 40 | 0.00 | 0.00 | 0.00 | 1.45 |
| Out_sounds | 13 | 0.71 | 0.11 | 4.41 | 0.71 | 13 | 0.60 | 0.19 | 0.91 | 0.73 | 13 | 0.13 | 0.02 | 1.00 | 0.79 | 13 | 0.16 | 0.02 | 0.61 | 0.83 |
| Out_sounds | 14 | 0.50 | 0.08 | 3.04 | 0.70 | 14 | 0.50 | 0.14 | 0.87 | 0.72 | 14 | 0.09 | 0.01 | 0.67 | 0.79 | 14 | 0.10 | 0.01 | 0.48 | 0.82 |
| Out_sounds | 15 | 0.35 | 0.06 | 2.10 | 0.70 | 15 | 0.40 | 0.10 | 0.81 | 0.71 | 15 | 0.06 | 0.01 | 0.45 | 0.78 | 15 | 0.07 | 0.01 | 0.37 | 0.81 |
| Out_sounds | 16 | 0.24 | 0.04 | 1.46 | 0.69 | 16 | 0.31 | 0.07 | 0.73 | 0.70 | 16 | 0.04 | 0.01 | 0.30 | 0.78 | 16 | 0.04 | 0.01 | 0.26 | 0.81 |
| Out_sounds | 17 | 0.17 | 0.03 | 1.02 | 0.69 | 17 | 0.23 | 0.05 | 0.64 | 0.69 | 17 | 0.03 | 0.00 | 0.21 | 0.78 | 17 | 0.03 | 0.00 | 0.18 | 0.82 |
| Out_sounds | 18 | 0.12 | 0.02 | 0.71 | 0.69 | 18 | 0.16 | 0.03 | 0.54 | 0.69 | 18 | 0.02 | 0.00 | 0.14 | 0.79 | 18 | 0.02 | 0.00 | 0.12 | 0.82 |
| Out_sounds | 19 | 0.08 | 0.01 | 0.50 | 0.70 | 19 | 0.12 | 0.02 | 0.44 | 0.70 | 19 | 0.01 | 0.00 | 0.10 | 0.79 | 19 | 0.01 | 0.00 | 0.08 | 0.83 |
| Out_sounds | 20 | 0.06 | 0.01 | 0.36 | 0.70 | 20 | 0.08 | 0.01 | 0.35 | 0.70 | 20 | 0.01 | 0.00 | 0.07 | 0.80 | 20 | 0.01 | 0.00 | 0.05 | 0.84 |
| Out_sounds | 21 | 0.04 | 0.01 | 0.25 | 0.71 | 21 | 0.05 | 0.01 | 0.27 | 0.71 | 21 | 0.01 | 0.00 | 0.05 | 0.80 | 21 | 0.00 | 0.00 | 0.04 | 0.86 |
| Out_sounds | 22 | 0.03 | 0.00 | 0.18 | 0.71 | 22 | 0.04 | 0.01 | 0.20 | 0.73 | 22 | 0.00 | 0.00 | 0.03 | 0.81 | 22 | 0.00 | 0.00 | 0.02 | 0.87 |
| Out_sounds | 23 | 0.02 | 0.00 | 0.13 | 0.72 | 23 | 0.02 | 0.00 | 0.15 | 0.75 | 23 | 0.00 | 0.00 | 0.02 | 0.82 | 23 | 0.00 | 0.00 | 0.02 | 0.89 |
| Out_sounds | 24 | 0.01 | 0.00 | 0.09 | 0.73 | 24 | 0.02 | 0.00 | 0.11 | 0.77 | 24 | 0.00 | 0.00 | 0.02 | 0.83 | 24 | 0.00 | 0.00 | 0.01 | 0.92 |
| Out_sounds | 25 | 0.01 | 0.00 | 0.07 | 0.75 | 25 | 0.01 | 0.00 | 0.08 | 0.79 | 25 | 0.00 | 0.00 | 0.01 | 0.85 | 25 | 0.00 | 0.00 | 0.01 | 0.94 |
| Out_sounds | 26 | 0.01 | 0.00 | 0.05 | 0.76 | 26 | 0.01 | 0.00 | 0.06 | 0.82 | 26 | 0.00 | 0.00 | 0.01 | 0.86 | 26 | 0.00 | 0.00 | 0.00 | 0.97 |
| Out_sounds | 27 | 0.00 | 0.00 | 0.04 | 0.78 | 27 | 0.00 | 0.00 | 0.04 | 0.84 | 27 | 0.00 | 0.00 | 0.01 | 0.88 | 27 | 0.00 | 0.00 | 0.00 | 1.00 |
| Out_sounds | 28 | 0.00 | 0.00 | 0.03 | 0.80 | 28 | 0.00 | 0.00 | 0.03 | 0.87 | 28 | 0.00 | 0.00 | 0.00 | 0.89 | 28 | 0.00 | 0.00 | 0.00 | 1.03 |
| Out_sounds | 29 | 0.00 | 0.00 | 0.02 | 0.81 | 29 | 0.00 | 0.00 | 0.02 | 0.91 | 29 | 0.00 | 0.00 | 0.00 | 0.91 | 29 | 0.00 | 0.00 | 0.00 | 1.06 |

| | | | | | | | | | | | | | | | | | | | | |
|---|---|---|---|---|---|---|---|---|---|---|---|---|---|---|---|---|---|---|---|---|
| Out_sounds | 30 | 0.00 | 0.00 | 0.01 | 0.83 | 30 | 0.00 | 0.00 | 0.02 | 0.94 | 30 | 0.00 | 0.00 | 0.00 | 0.93 | 30 | 0.00 | 0.00 | 0.00 | 1.10 |
| Out_sounds | 31 | 0.00 | 0.00 | 0.01 | 0.86 | 31 | 0.00 | 0.00 | 0.01 | 0.98 | 31 | 0.00 | 0.00 | 0.00 | 0.95 | 31 | 0.00 | 0.00 | 0.00 | 1.13 |
| Out_sounds | 32 | 0.00 | 0.00 | 0.01 | 0.88 | 32 | 0.00 | 0.00 | 0.01 | 1.01 | 32 | 0.00 | 0.00 | 0.00 | 0.97 | 32 | 0.00 | 0.00 | 0.00 | 1.17 |
| Out_sounds | 33 | 0.00 | 0.00 | 0.01 | 0.90 | 33 | 0.00 | 0.00 | 0.01 | 1.05 | 33 | 0.00 | 0.00 | 0.00 | 0.99 | 33 | 0.00 | 0.00 | 0.00 | 1.21 |
| Out_sounds | 34 | 0.00 | 0.00 | 0.00 | 0.92 | 34 | 0.00 | 0.00 | 0.00 | 1.09 | 34 | 0.00 | 0.00 | 0.00 | 1.01 | 34 | 0.00 | 0.00 | 0.00 | 1.25 |
| Out_sounds | 35 | 0.00 | 0.00 | 0.00 | 0.95 | 35 | 0.00 | 0.00 | 0.00 | 1.13 | 35 | 0.00 | 0.00 | 0.00 | 1.03 | 35 | 0.00 | 0.00 | 0.00 | 1.29 |
| Out_sounds | 36 | 0.00 | 0.00 | 0.00 | 0.97 | 36 | 0.00 | 0.00 | 0.00 | 1.18 | 36 | 0.00 | 0.00 | 0.00 | 1.06 | 36 | 0.00 | 0.00 | 0.00 | 1.33 |
| Out_sounds | 37 | 0.00 | 0.00 | 0.00 | 1.00 | 37 | 0.00 | 0.00 | 0.00 | 1.22 | 37 | 0.00 | 0.00 | 0.00 | 1.08 | 37 | 0.00 | 0.00 | 0.00 | 1.37 |
| Out_sounds | 38 | 0.00 | 0.00 | 0.00 | 1.03 | 38 | 0.00 | 0.00 | 0.00 | 1.26 | 38 | 0.00 | 0.00 | 0.00 | 1.11 | 38 | 0.00 | 0.00 | 0.00 | 1.41 |
| Out_sounds | 39 | 0.00 | 0.00 | 0.00 | 1.06 | 39 | 0.00 | 0.00 | 0.00 | 1.30 | 39 | 0.00 | 0.00 | 0.00 | 1.13 | 39 | 0.00 | 0.00 | 0.00 | 1.45 |
| Out_sounds | 40 | 0.00 | 0.00 | 0.00 | 1.08 | 40 | 0.00 | 0.00 | 0.00 | 1.35 | 40 | 0.00 | 0.00 | 0.00 | 1.16 | 40 | 0.00 | 0.00 | 0.00 | 1.50 |
| Pel_back | 13 | 0.44 | 0.08 | 2.30 | 0.64 | 13 | 0.46 | 0.15 | 0.81 | 0.62 | 13 | 0.10 | 0.02 | 0.64 | 0.71 | 13 | 0.12 | 0.02 | 0.49 | 0.75 |
| Pel_back | 14 | 0.31 | 0.06 | 1.59 | 0.64 | 14 | 0.37 | 0.11 | 0.73 | 0.60 | 14 | 0.07 | 0.01 | 0.43 | 0.71 | 14 | 0.08 | 0.01 | 0.37 | 0.74 |
| Pel_back | 15 | 0.22 | 0.04 | 1.10 | 0.63 | 15 | 0.28 | 0.08 | 0.64 | 0.59 | 15 | 0.05 | 0.01 | 0.29 | 0.70 | 15 | 0.05 | 0.01 | 0.27 | 0.74 |
| Pel_back | 16 | 0.15 | 0.03 | 0.77 | 0.63 | 16 | 0.20 | 0.05 | 0.54 | 0.59 | 16 | 0.03 | 0.01 | 0.20 | 0.71 | 16 | 0.03 | 0.00 | 0.18 | 0.74 |
| Pel_back | 17 | 0.11 | 0.02 | 0.54 | 0.63 | 17 | 0.14 | 0.04 | 0.43 | 0.59 | 17 | 0.02 | 0.00 | 0.13 | 0.71 | 17 | 0.02 | 0.00 | 0.13 | 0.75 |
| Pel_back | 18 | 0.07 | 0.01 | 0.38 | 0.63 | 18 | 0.10 | 0.02 | 0.34 | 0.59 | 18 | 0.01 | 0.00 | 0.09 | 0.71 | 18 | 0.01 | 0.00 | 0.08 | 0.75 |
| Pel_back | 19 | 0.05 | 0.01 | 0.27 | 0.64 | 19 | 0.07 | 0.02 | 0.26 | 0.60 | 19 | 0.01 | 0.00 | 0.06 | 0.72 | 19 | 0.01 | 0.00 | 0.06 | 0.77 |
| Pel_back | 20 | 0.04 | 0.01 | 0.19 | 0.64 | 20 | 0.05 | 0.01 | 0.19 | 0.61 | 20 | 0.01 | 0.00 | 0.04 | 0.73 | 20 | 0.01 | 0.00 | 0.04 | 0.78 |
| Pel_back | 21 | 0.03 | 0.00 | 0.14 | 0.65 | 21 | 0.03 | 0.01 | 0.14 | 0.63 | 21 | 0.00 | 0.00 | 0.03 | 0.73 | 21 | 0.00 | 0.00 | 0.02 | 0.80 |
| Pel_back | 22 | 0.02 | 0.00 | 0.10 | 0.66 | 22 | 0.02 | 0.00 | 0.10 | 0.65 | 22 | 0.00 | 0.00 | 0.02 | 0.75 | 22 | 0.00 | 0.00 | 0.02 | 0.82 |
| Pel_back | 23 | 0.01 | 0.00 | 0.07 | 0.67 | 23 | 0.01 | 0.00 | 0.08 | 0.67 | 23 | 0.00 | 0.00 | 0.01 | 0.76 | 23 | 0.00 | 0.00 | 0.01 | 0.85 |
| Pel_back | 24 | 0.01 | 0.00 | 0.05 | 0.69 | 24 | 0.01 | 0.00 | 0.05 | 0.70 | 24 | 0.00 | 0.00 | 0.01 | 0.77 | 24 | 0.00 | 0.00 | 0.01 | 0.87 |
| Pel_back | 25 | 0.01 | 0.00 | 0.04 | 0.70 | 25 | 0.01 | 0.00 | 0.04 | 0.72 | 25 | 0.00 | 0.00 | 0.01 | 0.79 | 25 | 0.00 | 0.00 | 0.00 | 0.90 |
| Pel_back | 26 | 0.00 | 0.00 | 0.03 | 0.72 | 26 | 0.00 | 0.00 | 0.03 | 0.76 | 26 | 0.00 | 0.00 | 0.01 | 0.80 | 26 | 0.00 | 0.00 | 0.00 | 0.93 |
| Pel_back | 27 | 0.00 | 0.00 | 0.02 | 0.74 | 27 | 0.00 | 0.00 | 0.02 | 0.79 | 27 | 0.00 | 0.00 | 0.00 | 0.82 | 27 | 0.00 | 0.00 | 0.00 | 0.96 |
| Pel_back | 28 | 0.00 | 0.00 | 0.02 | 0.76 | 28 | 0.00 | 0.00 | 0.02 | 0.82 | 28 | 0.00 | 0.00 | 0.00 | 0.84 | 28 | 0.00 | 0.00 | 0.00 | 1.00 |
| Pel_back | 29 | 0.00 | 0.00 | 0.01 | 0.78 | 29 | 0.00 | 0.00 | 0.01 | 0.86 | 29 | 0.00 | 0.00 | 0.00 | 0.86 | 29 | 0.00 | 0.00 | 0.00 | 1.03 |
| Pel_back | 30 | 0.00 | 0.00 | 0.01 | 0.80 | 30 | 0.00 | 0.00 | 0.01 | 0.90 | 30 | 0.00 | 0.00 | 0.00 | 0.88 | 30 | 0.00 | 0.00 | 0.00 | 1.07 |
| Pel_back | 31 | 0.00 | 0.00 | 0.01 | 0.82 | 31 | 0.00 | 0.00 | 0.01 | 0.94 | 31 | 0.00 | 0.00 | 0.00 | 0.90 | 31 | 0.00 | 0.00 | 0.00 | 1.11 |
| Pel_back | 32 | 0.00 | 0.00 | 0.00 | 0.85 | 32 | 0.00 | 0.00 | 0.00 | 0.98 | 32 | 0.00 | 0.00 | 0.00 | 0.93 | 32 | 0.00 | 0.00 | 0.00 | 1.15 |

| | | | | | | | | | | | | | | | | | | | | |
|---|---|---|---|---|---|---|---|---|---|---|---|---|---|---|---|---|---|---|---|---|
| Pel_back | 33 | 0.00 | 0.00 | 0.00 | 0.87 | 33 | 0.00 | 0.00 | 0.00 | 1.02 | 33 | 0.00 | 0.00 | 0.00 | 0.95 | 33 | 0.00 | 0.00 | 0.00 | 1.19 |
| Pel_back | 34 | 0.00 | 0.00 | 0.00 | 0.90 | 34 | 0.00 | 0.00 | 0.00 | 1.07 | 34 | 0.00 | 0.00 | 0.00 | 0.97 | 34 | 0.00 | 0.00 | 0.00 | 1.23 |
| Pel_back | 35 | 0.00 | 0.00 | 0.00 | 0.93 | 35 | 0.00 | 0.00 | 0.00 | 1.11 | 35 | 0.00 | 0.00 | 0.00 | 1.00 | 35 | 0.00 | 0.00 | 0.00 | 1.27 |
| Pel_back | 36 | 0.00 | 0.00 | 0.00 | 0.95 | 36 | 0.00 | 0.00 | 0.00 | 1.15 | 36 | 0.00 | 0.00 | 0.00 | 1.02 | 36 | 0.00 | 0.00 | 0.00 | 1.31 |
| Pel_back | 37 | 0.00 | 0.00 | 0.00 | 0.98 | 37 | 0.00 | 0.00 | 0.00 | 1.20 | 37 | 0.00 | 0.00 | 0.00 | 1.05 | 37 | 0.00 | 0.00 | 0.00 | 1.36 |
| Pel_back | 38 | 0.00 | 0.00 | 0.00 | 1.01 | 38 | 0.00 | 0.00 | 0.00 | 1.24 | 38 | 0.00 | 0.00 | 0.00 | 1.08 | 38 | 0.00 | 0.00 | 0.00 | 1.40 |
| Pel_back | 39 | 0.00 | 0.00 | 0.00 | 1.04 | 39 | 0.00 | 0.00 | 0.00 | 1.29 | 39 | 0.00 | 0.00 | 0.00 | 1.10 | 39 | 0.00 | 0.00 | 0.00 | 1.45 |
| Pel_back | 40 | 0.00 | 0.00 | 0.00 | 1.07 | 40 | 0.00 | 0.00 | 0.00 | 1.34 | 40 | 0.00 | 0.00 | 0.00 | 1.13 | 40 | 0.00 | 0.00 | 0.00 | 1.49 |
| Pel_central | 13 | 1.42 | 0.51 | 3.98 | 0.40 | 13 | 0.57 | 0.27 | 0.82 | 0.49 | 13 | 0.96 | 0.24 | 3.79 | 0.53 | 13 | 0.47 | 0.14 | 0.83 | 0.66 |
| Pel_central | 14 | 1.00 | 0.36 | 2.73 | 0.39 | 14 | 0.47 | 0.20 | 0.75 | 0.48 | 14 | 0.65 | 0.17 | 2.52 | 0.53 | 14 | 0.36 | 0.09 | 0.75 | 0.65 |
| Pel_central | 15 | 0.70 | 0.26 | 1.89 | 0.39 | 15 | 0.37 | 0.15 | 0.66 | 0.47 | 15 | 0.44 | 0.11 | 1.69 | 0.52 | 15 | 0.26 | 0.06 | 0.65 | 0.64 |
| Pel_central | 16 | 0.49 | 0.18 | 1.32 | 0.39 | 16 | 0.28 | 0.10 | 0.56 | 0.46 | 16 | 0.30 | 0.08 | 1.14 | 0.52 | 16 | 0.18 | 0.04 | 0.53 | 0.64 |
| Pel_central | 17 | 0.34 | 0.13 | 0.93 | 0.39 | 17 | 0.20 | 0.07 | 0.46 | 0.47 | 17 | 0.20 | 0.05 | 0.77 | 0.52 | 17 | 0.12 | 0.03 | 0.41 | 0.64 |
| Pel_central | 18 | 0.24 | 0.09 | 0.66 | 0.39 | 18 | 0.15 | 0.05 | 0.37 | 0.48 | 18 | 0.14 | 0.03 | 0.53 | 0.53 | 18 | 0.08 | 0.02 | 0.31 | 0.65 |
| Pel_central | 19 | 0.17 | 0.06 | 0.47 | 0.40 | 19 | 0.10 | 0.03 | 0.28 | 0.49 | 19 | 0.09 | 0.02 | 0.36 | 0.53 | 19 | 0.05 | 0.01 | 0.22 | 0.66 |
| Pel_central | 20 | 0.12 | 0.04 | 0.34 | 0.41 | 20 | 0.07 | 0.02 | 0.22 | 0.51 | 20 | 0.06 | 0.02 | 0.25 | 0.54 | 20 | 0.03 | 0.01 | 0.15 | 0.67 |
| Pel_central | 21 | 0.08 | 0.03 | 0.25 | 0.43 | 21 | 0.05 | 0.01 | 0.16 | 0.53 | 21 | 0.04 | 0.01 | 0.17 | 0.55 | 21 | 0.02 | 0.00 | 0.11 | 0.69 |
| Pel_central | 22 | 0.06 | 0.02 | 0.18 | 0.44 | 22 | 0.03 | 0.01 | 0.12 | 0.56 | 22 | 0.03 | 0.01 | 0.12 | 0.56 | 22 | 0.01 | 0.00 | 0.07 | 0.71 |
| Pel_central | 23 | 0.04 | 0.01 | 0.13 | 0.46 | 23 | 0.02 | 0.00 | 0.09 | 0.59 | 23 | 0.02 | 0.00 | 0.09 | 0.58 | 23 | 0.01 | 0.00 | 0.05 | 0.73 |
| Pel_central | 24 | 0.03 | 0.01 | 0.10 | 0.48 | 24 | 0.01 | 0.00 | 0.07 | 0.62 | 24 | 0.01 | 0.00 | 0.06 | 0.59 | 24 | 0.00 | 0.00 | 0.03 | 0.76 |
| Pel_central | 25 | 0.02 | 0.01 | 0.07 | 0.51 | 25 | 0.01 | 0.00 | 0.05 | 0.66 | 25 | 0.01 | 0.00 | 0.04 | 0.61 | 25 | 0.00 | 0.00 | 0.02 | 0.79 |
| Pel_central | 26 | 0.01 | 0.00 | 0.06 | 0.53 | 26 | 0.01 | 0.00 | 0.04 | 0.69 | 26 | 0.01 | 0.00 | 0.03 | 0.63 | 26 | 0.00 | 0.00 | 0.02 | 0.82 |
| Pel_central | 27 | 0.01 | 0.00 | 0.04 | 0.56 | 27 | 0.00 | 0.00 | 0.03 | 0.73 | 27 | 0.00 | 0.00 | 0.02 | 0.65 | 27 | 0.00 | 0.00 | 0.01 | 0.85 |
| Pel_central | 28 | 0.01 | 0.00 | 0.03 | 0.59 | 28 | 0.00 | 0.00 | 0.02 | 0.77 | 28 | 0.00 | 0.00 | 0.02 | 0.67 | 28 | 0.00 | 0.00 | 0.01 | 0.89 |
| Pel_central | 29 | 0.00 | 0.00 | 0.02 | 0.61 | 29 | 0.00 | 0.00 | 0.02 | 0.82 | 29 | 0.00 | 0.00 | 0.01 | 0.70 | 29 | 0.00 | 0.00 | 0.00 | 0.92 |
| Pel_central | 30 | 0.00 | 0.00 | 0.02 | 0.64 | 30 | 0.00 | 0.00 | 0.01 | 0.86 | 30 | 0.00 | 0.00 | 0.01 | 0.72 | 30 | 0.00 | 0.00 | 0.00 | 0.96 |
| Pel_central | 31 | 0.00 | 0.00 | 0.01 | 0.67 | 31 | 0.00 | 0.00 | 0.01 | 0.90 | 31 | 0.00 | 0.00 | 0.01 | 0.74 | 31 | 0.00 | 0.00 | 0.00 | 1.00 |
| Pel_central | 32 | 0.00 | 0.00 | 0.01 | 0.70 | 32 | 0.00 | 0.00 | 0.01 | 0.95 | 32 | 0.00 | 0.00 | 0.00 | 0.77 | 32 | 0.00 | 0.00 | 0.00 | 1.04 |
| Pel_central | 33 | 0.00 | 0.00 | 0.01 | 0.73 | 33 | 0.00 | 0.00 | 0.00 | 0.99 | 33 | 0.00 | 0.00 | 0.00 | 0.80 | 33 | 0.00 | 0.00 | 0.00 | 1.08 |
| Pel_central | 34 | 0.00 | 0.00 | 0.01 | 0.76 | 34 | 0.00 | 0.00 | 0.00 | 1.04 | 34 | 0.00 | 0.00 | 0.00 | 0.82 | 34 | 0.00 | 0.00 | 0.00 | 1.12 |
| Pel_central | 35 | 0.00 | 0.00 | 0.00 | 0.80 | 35 | 0.00 | 0.00 | 0.00 | 1.09 | 35 | 0.00 | 0.00 | 0.00 | 0.85 | 35 | 0.00 | 0.00 | 0.00 | 1.17 |

| | | | | | | | | | | | | | | | | | | | | |
|---|---|---|---|---|---|---|---|---|---|---|---|---|---|---|---|---|---|---|---|---|
| Pel_central | 36 | 0.00 | 0.00 | 0.00 | 0.83 | 36 | 0.00 | 0.00 | 0.00 | 1.13 | 36 | 0.00 | 0.00 | 0.00 | 0.88 | 36 | 0.00 | 0.00 | 0.00 | 1.21 |
| Pel_central | 37 | 0.00 | 0.00 | 0.00 | 0.86 | 37 | 0.00 | 0.00 | 0.00 | 1.18 | 37 | 0.00 | 0.00 | 0.00 | 0.91 | 37 | 0.00 | 0.00 | 0.00 | 1.26 |
| Pel_central | 38 | 0.00 | 0.00 | 0.00 | 0.89 | 38 | 0.00 | 0.00 | 0.00 | 1.23 | 38 | 0.00 | 0.00 | 0.00 | 0.94 | 38 | 0.00 | 0.00 | 0.00 | 1.30 |
| Pel_central | 39 | 0.00 | 0.00 | 0.00 | 0.93 | 39 | 0.00 | 0.00 | 0.00 | 1.28 | 39 | 0.00 | 0.00 | 0.00 | 0.97 | 39 | 0.00 | 0.00 | 0.00 | 1.35 |
| Pel_central | 40 | 0.00 | 0.00 | 0.00 | 0.96 | 40 | 0.00 | 0.00 | 0.00 | 1.32 | 40 | 0.00 | 0.00 | 0.00 | 1.00 | 40 | 0.00 | 0.00 | 0.00 | 1.39 |
| Pelorus | 13 | 0.86 | 0.28 | 2.63 | 0.43 | 13 | 0.63 | 0.35 | 0.84 | 0.44 | 13 | 0.81 | 0.18 | 3.57 | 0.58 | 13 | 0.63 | 0.23 | 0.91 | 0.67 |
| Pelorus | 14 | 0.60 | 0.20 | 1.81 | 0.43 | 14 | 0.53 | 0.27 | 0.77 | 0.43 | 14 | 0.55 | 0.13 | 2.39 | 0.57 | 14 | 0.52 | 0.16 | 0.85 | 0.66 |
| Pelorus | 15 | 0.42 | 0.14 | 1.26 | 0.42 | 15 | 0.43 | 0.20 | 0.68 | 0.42 | 15 | 0.37 | 0.09 | 1.61 | 0.57 | 15 | 0.40 | 0.11 | 0.78 | 0.65 |
| Pelorus | 16 | 0.30 | 0.10 | 0.88 | 0.42 | 16 | 0.33 | 0.15 | 0.59 | 0.41 | 16 | 0.25 | 0.06 | 1.09 | 0.57 | 16 | 0.29 | 0.07 | 0.69 | 0.65 |
| Pelorus | 17 | 0.21 | 0.07 | 0.63 | 0.43 | 17 | 0.25 | 0.10 | 0.49 | 0.41 | 17 | 0.17 | 0.04 | 0.74 | 0.57 | 17 | 0.20 | 0.05 | 0.58 | 0.65 |
| Pelorus | 18 | 0.15 | 0.05 | 0.45 | 0.43 | 18 | 0.18 | 0.07 | 0.39 | 0.42 | 18 | 0.11 | 0.03 | 0.51 | 0.58 | 18 | 0.14 | 0.03 | 0.46 | 0.65 |
| Pelorus | 19 | 0.10 | 0.03 | 0.32 | 0.44 | 19 | 0.13 | 0.05 | 0.31 | 0.44 | 19 | 0.08 | 0.02 | 0.35 | 0.58 | 19 | 0.09 | 0.02 | 0.35 | 0.66 |
| Pelorus | 20 | 0.07 | 0.02 | 0.23 | 0.46 | 20 | 0.09 | 0.03 | 0.24 | 0.46 | 20 | 0.05 | 0.01 | 0.24 | 0.59 | 20 | 0.06 | 0.01 | 0.26 | 0.67 |
| Pelorus | 21 | 0.05 | 0.01 | 0.17 | 0.47 | 21 | 0.06 | 0.02 | 0.18 | 0.48 | 21 | 0.04 | 0.01 | 0.17 | 0.60 | 21 | 0.04 | 0.01 | 0.19 | 0.69 |
| Pelorus | 22 | 0.04 | 0.01 | 0.12 | 0.49 | 22 | 0.04 | 0.01 | 0.14 | 0.51 | 22 | 0.02 | 0.00 | 0.12 | 0.61 | 22 | 0.02 | 0.00 | 0.13 | 0.71 |
| Pelorus | 23 | 0.02 | 0.01 | 0.09 | 0.51 | 23 | 0.03 | 0.01 | 0.10 | 0.54 | 23 | 0.02 | 0.00 | 0.08 | 0.63 | 23 | 0.01 | 0.00 | 0.09 | 0.73 |
| Pelorus | 24 | 0.02 | 0.00 | 0.07 | 0.53 | 24 | 0.02 | 0.00 | 0.08 | 0.58 | 24 | 0.01 | 0.00 | 0.06 | 0.64 | 24 | 0.01 | 0.00 | 0.06 | 0.76 |
| Pelorus | 25 | 0.01 | 0.00 | 0.05 | 0.55 | 25 | 0.01 | 0.00 | 0.06 | 0.61 | 25 | 0.01 | 0.00 | 0.04 | 0.66 | 25 | 0.01 | 0.00 | 0.04 | 0.79 |
| Pelorus | 26 | 0.01 | 0.00 | 0.04 | 0.57 | 26 | 0.01 | 0.00 | 0.04 | 0.65 | 26 | 0.01 | 0.00 | 0.03 | 0.68 | 26 | 0.00 | 0.00 | 0.03 | 0.82 |
| Pelorus | 27 | 0.01 | 0.00 | 0.03 | 0.60 | 27 | 0.01 | 0.00 | 0.03 | 0.69 | 27 | 0.00 | 0.00 | 0.02 | 0.70 | 27 | 0.00 | 0.00 | 0.02 | 0.85 |
| Pelorus | 28 | 0.00 | 0.00 | 0.02 | 0.63 | 28 | 0.00 | 0.00 | 0.02 | 0.74 | 28 | 0.00 | 0.00 | 0.01 | 0.72 | 28 | 0.00 | 0.00 | 0.01 | 0.88 |
| Pelorus | 29 | 0.00 | 0.00 | 0.02 | 0.65 | 29 | 0.00 | 0.00 | 0.02 | 0.78 | 29 | 0.00 | 0.00 | 0.01 | 0.74 | 29 | 0.00 | 0.00 | 0.01 | 0.92 |
| Pelorus | 30 | 0.00 | 0.00 | 0.01 | 0.68 | 30 | 0.00 | 0.00 | 0.01 | 0.82 | 30 | 0.00 | 0.00 | 0.01 | 0.77 | 30 | 0.00 | 0.00 | 0.01 | 0.96 |
| Pelorus | 31 | 0.00 | 0.00 | 0.01 | 0.71 | 31 | 0.00 | 0.00 | 0.01 | 0.87 | 31 | 0.00 | 0.00 | 0.01 | 0.79 | 31 | 0.00 | 0.00 | 0.00 | 1.00 |
| Pelorus | 32 | 0.00 | 0.00 | 0.01 | 0.74 | 32 | 0.00 | 0.00 | 0.01 | 0.91 | 32 | 0.00 | 0.00 | 0.00 | 0.82 | 32 | 0.00 | 0.00 | 0.00 | 1.04 |
| Pelorus | 33 | 0.00 | 0.00 | 0.01 | 0.77 | 33 | 0.00 | 0.00 | 0.01 | 0.96 | 33 | 0.00 | 0.00 | 0.00 | 0.84 | 33 | 0.00 | 0.00 | 0.00 | 1.08 |
| Pelorus | 34 | 0.00 | 0.00 | 0.00 | 0.80 | 34 | 0.00 | 0.00 | 0.00 | 1.01 | 34 | 0.00 | 0.00 | 0.00 | 0.87 | 34 | 0.00 | 0.00 | 0.00 | 1.12 |
| Pelorus | 35 | 0.00 | 0.00 | 0.00 | 0.83 | 35 | 0.00 | 0.00 | 0.00 | 1.06 | 35 | 0.00 | 0.00 | 0.00 | 0.90 | 35 | 0.00 | 0.00 | 0.00 | 1.16 |
| Pelorus | 36 | 0.00 | 0.00 | 0.00 | 0.86 | 36 | 0.00 | 0.00 | 0.00 | 1.10 | 36 | 0.00 | 0.00 | 0.00 | 0.93 | 36 | 0.00 | 0.00 | 0.00 | 1.20 |
| Pelorus | 37 | 0.00 | 0.00 | 0.00 | 0.90 | 37 | 0.00 | 0.00 | 0.00 | 1.15 | 37 | 0.00 | 0.00 | 0.00 | 0.95 | 37 | 0.00 | 0.00 | 0.00 | 1.25 |
| Pelorus | 38 | 0.00 | 0.00 | 0.00 | 0.93 | 38 | 0.00 | 0.00 | 0.00 | 1.20 | 38 | 0.00 | 0.00 | 0.00 | 0.98 | 38 | 0.00 | 0.00 | 0.00 | 1.29 |

| | | | | | | | | | | | | | | | | | | | | |
|---|---|---|---|---|---|---|---|---|---|---|---|---|---|---|---|---|---|---|---|---|
| Pelorus | 39 | 0.00 | 0.00 | 0.00 | 0.96 | 39 | 0.00 | 0.00 | 0.00 | 1.25 | 39 | 0.00 | 0.00 | 0.00 | 1.01 | 39 | 0.00 | 0.00 | 0.00 | 1.34 |
| Pelorus | 40 | 0.00 | 0.00 | 0.00 | 1.00 | 40 | 0.00 | 0.00 | 0.00 | 1.30 | 40 | 0.00 | 0.00 | 0.00 | 1.04 | 40 | 0.00 | 0.00 | 0.00 | 1.38 |
| Tennyson | 13 | 0.11 | 0.01 | 2.15 | 1.15 | 13 | 0.14 | 0.01 | 0.73 | 1.08 | 13 | 0.03 | 0.00 | 0.92 | 1.29 | 13 | 0.04 | 0.00 | 0.54 | 1.32 |
| Tennyson | 14 | 0.08 | 0.00 | 1.50 | 1.15 | 14 | 0.10 | 0.01 | 0.64 | 1.08 | 14 | 0.02 | 0.00 | 0.62 | 1.29 | 14 | 0.02 | 0.00 | 0.42 | 1.32 |
| Tennyson | 15 | 0.05 | 0.00 | 1.05 | 1.15 | 15 | 0.07 | 0.00 | 0.54 | 1.08 | 15 | 0.02 | 0.00 | 0.42 | 1.29 | 15 | 0.01 | 0.00 | 0.31 | 1.32 |
| Tennyson | 16 | 0.04 | 0.00 | 0.74 | 1.15 | 16 | 0.05 | 0.00 | 0.44 | 1.08 | 16 | 0.01 | 0.00 | 0.28 | 1.29 | 16 | 0.01 | 0.00 | 0.22 | 1.32 |
| Tennyson | 17 | 0.03 | 0.00 | 0.52 | 1.16 | 17 | 0.03 | 0.00 | 0.35 | 1.09 | 17 | 0.01 | 0.00 | 0.19 | 1.29 | 17 | 0.01 | 0.00 | 0.15 | 1.33 |
| Tennyson | 18 | 0.02 | 0.00 | 0.37 | 1.16 | 18 | 0.02 | 0.00 | 0.26 | 1.09 | 18 | 0.00 | 0.00 | 0.13 | 1.29 | 18 | 0.00 | 0.00 | 0.10 | 1.33 |
| Tennyson | 19 | 0.01 | 0.00 | 0.26 | 1.16 | 19 | 0.01 | 0.00 | 0.20 | 1.10 | 19 | 0.00 | 0.00 | 0.09 | 1.29 | 19 | 0.00 | 0.00 | 0.07 | 1.34 |
| Tennyson | 20 | 0.01 | 0.00 | 0.19 | 1.17 | 20 | 0.01 | 0.00 | 0.14 | 1.11 | 20 | 0.00 | 0.00 | 0.06 | 1.30 | 20 | 0.00 | 0.00 | 0.04 | 1.35 |
| Tennyson | 21 | 0.01 | 0.00 | 0.13 | 1.17 | 21 | 0.01 | 0.00 | 0.10 | 1.13 | 21 | 0.00 | 0.00 | 0.04 | 1.30 | 21 | 0.00 | 0.00 | 0.03 | 1.36 |
| Tennyson | 22 | 0.00 | 0.00 | 0.09 | 1.18 | 22 | 0.00 | 0.00 | 0.07 | 1.14 | 22 | 0.00 | 0.00 | 0.03 | 1.31 | 22 | 0.00 | 0.00 | 0.02 | 1.37 |
| Tennyson | 23 | 0.00 | 0.00 | 0.07 | 1.19 | 23 | 0.00 | 0.00 | 0.05 | 1.16 | 23 | 0.00 | 0.00 | 0.02 | 1.32 | 23 | 0.00 | 0.00 | 0.01 | 1.39 |
| Tennyson | 24 | 0.00 | 0.00 | 0.05 | 1.20 | 24 | 0.00 | 0.00 | 0.04 | 1.18 | 24 | 0.00 | 0.00 | 0.01 | 1.33 | 24 | 0.00 | 0.00 | 0.01 | 1.41 |
| Tennyson | 25 | 0.00 | 0.00 | 0.04 | 1.21 | 25 | 0.00 | 0.00 | 0.03 | 1.20 | 25 | 0.00 | 0.00 | 0.01 | 1.33 | 25 | 0.00 | 0.00 | 0.01 | 1.42 |
| Tennyson | 26 | 0.00 | 0.00 | 0.03 | 1.22 | 26 | 0.00 | 0.00 | 0.02 | 1.23 | 26 | 0.00 | 0.00 | 0.01 | 1.34 | 26 | 0.00 | 0.00 | 0.00 | 1.45 |
| Tennyson | 27 | 0.00 | 0.00 | 0.02 | 1.24 | 27 | 0.00 | 0.00 | 0.01 | 1.25 | 27 | 0.00 | 0.00 | 0.00 | 1.36 | 27 | 0.00 | 0.00 | 0.00 | 1.47 |
| Tennyson | 28 | 0.00 | 0.00 | 0.01 | 1.25 | 28 | 0.00 | 0.00 | 0.01 | 1.28 | 28 | 0.00 | 0.00 | 0.00 | 1.37 | 28 | 0.00 | 0.00 | 0.00 | 1.49 |
| Tennyson | 29 | 0.00 | 0.00 | 0.01 | 1.26 | 29 | 0.00 | 0.00 | 0.01 | 1.31 | 29 | 0.00 | 0.00 | 0.00 | 1.38 | 29 | 0.00 | 0.00 | 0.00 | 1.52 |
| Tennyson | 30 | 0.00 | 0.00 | 0.01 | 1.28 | 30 | 0.00 | 0.00 | 0.00 | 1.34 | 30 | 0.00 | 0.00 | 0.00 | 1.39 | 30 | 0.00 | 0.00 | 0.00 | 1.54 |
| Tennyson | 31 | 0.00 | 0.00 | 0.01 | 1.30 | 31 | 0.00 | 0.00 | 0.00 | 1.37 | 31 | 0.00 | 0.00 | 0.00 | 1.41 | 31 | 0.00 | 0.00 | 0.00 | 1.57 |
| Tennyson | 32 | 0.00 | 0.00 | 0.00 | 1.31 | 32 | 0.00 | 0.00 | 0.00 | 1.40 | 32 | 0.00 | 0.00 | 0.00 | 1.42 | 32 | 0.00 | 0.00 | 0.00 | 1.60 |
| Tennyson | 33 | 0.00 | 0.00 | 0.00 | 1.33 | 33 | 0.00 | 0.00 | 0.00 | 1.43 | 33 | 0.00 | 0.00 | 0.00 | 1.44 | 33 | 0.00 | 0.00 | 0.00 | 1.63 |
| Tennyson | 34 | 0.00 | 0.00 | 0.00 | 1.35 | 34 | 0.00 | 0.00 | 0.00 | 1.47 | 34 | 0.00 | 0.00 | 0.00 | 1.45 | 34 | 0.00 | 0.00 | 0.00 | 1.66 |
| Tennyson | 35 | 0.00 | 0.00 | 0.00 | 1.37 | 35 | 0.00 | 0.00 | 0.00 | 1.50 | 35 | 0.00 | 0.00 | 0.00 | 1.47 | 35 | 0.00 | 0.00 | 0.00 | 1.69 |
| Tennyson | 36 | 0.00 | 0.00 | 0.00 | 1.39 | 36 | 0.00 | 0.00 | 0.00 | 1.54 | 36 | 0.00 | 0.00 | 0.00 | 1.49 | 36 | 0.00 | 0.00 | 0.00 | 1.72 |
| Tennyson | 37 | 0.00 | 0.00 | 0.00 | 1.41 | 37 | 0.00 | 0.00 | 0.00 | 1.58 | 37 | 0.00 | 0.00 | 0.00 | 1.51 | 37 | 0.00 | 0.00 | 0.00 | 1.76 |
| Tennyson | 38 | 0.00 | 0.00 | 0.00 | 1.43 | 38 | 0.00 | 0.00 | 0.00 | 1.62 | 38 | 0.00 | 0.00 | 0.00 | 1.53 | 38 | 0.00 | 0.00 | 0.00 | 1.79 |
| Tennyson | 39 | 0.00 | 0.00 | 0.00 | 1.45 | 39 | 0.00 | 0.00 | 0.00 | 1.65 | 39 | 0.00 | 0.00 | 0.00 | 1.55 | 39 | 0.00 | 0.00 | 0.00 | 1.83 |
| Tennyson | 40 | 0.00 | 0.00 | 0.00 | 1.48 | 40 | 0.00 | 0.00 | 0.00 | 1.69 | 40 | 0.00 | 0.00 | 0.00 | 1.57 | 40 | 0.00 | 0.00 | 0.00 | 1.86 |
